# PIP4K attenuates PIP5K lipid kinase activity by disrupting membrane-mediated dimerization

**DOI:** 10.1101/2025.09.26.678889

**Authors:** Benjamin R. Duewell, Michael J. Chirumbolo, Samantha M. Fernandez-Ortiz, Gerald R.V. Hammond, Scott D. Hansen

## Abstract

The phosphatidylinositol 4-phosphate 5-kinase (PIP5K) family of enzymes generate most of the phosphatidylinositol-4,5-bisphosphate (PI(4,5)P_2_) lipids in eukaryotes. In solution, PIP5K exists in a weak monomer-dimer equilibrium but undergoes membrane-mediated dimerization, which potentiates lipid kinase activity. In vivo, however, PI(4,5)P_2_ levels are held remarkably constant due to the homeostatic regulation of PIP5K by PIP4K. We hypothesized that mechanisms that regulate PIP5K dimerization could function to buffer lipid kinase activity, thus providing a mechanism for maintaining relatively constant PI(4,5)P_2_ levels at the plasma membrane. Due the transient nature and density dependence of PIP5K dimerization, deciphering how other proteins modulate PIP5K dimerization has not been feasible. To address this limitation, we established a single molecule FRET assay to visualize membrane-mediated homodimerization and heterodimerization of PIP5K paralogs on supported lipid bilayers using Total Internal Reflection Fluorescence Microscopy (TIRF-M). Using this approach, we find that PIP4K attenuates PIP5K lipid kinase activity by disrupting membrane-mediated dimerization. Guided by structure prediction, we generated PIP4K mutants that are unable to disrupt PIP5K membrane-mediated dimerization thus preventing the attenuation lipid kinase activity. In vivo, mutations that disrupt the PIP4K-PIP5K interaction similarly prevent PIP4K-mediated inhibition of the PIP5K activity. Overall, this work reveals the molecular basis of the PIP4K-mediated inhibition of PIP5K, which underlies PI(4,5)P_2_ lipid homeostasis. Creation of this PIP5K dimerization FRET biosensor also establishes a novel tool for deciphering how proteins modulate membrane-mediated dimerization of PIP5K in the future.

## INTRODUCTION

Despite being only a minor component of eukaryotic cell membranes, phosphatidylinositol-4,5-bisphosphate (PI(4,5)P_2_) plays a major functional role in membrane signaling pathways (1, 2). PI(4,5)P_2_ function as lipid cofactors that recruit numerous peripheral membrane binding proteins to the inner leaflet of the plasma membrane, and also allosterically active integral and peripheral membrane proteins. This cofactor role has been shown to be critical for ion channel gating, cell signaling, membrane-cytoskeletal anchoring, and endocytosis (3–9). PI(4,5)P_2_ lipids also function as a precursor for the production of PI(3,4,5)P_3_ lipids, which controls cell fate decisions, directed cell migration, neurogenesis, and adhesion (10–12). These key roles make PI(4,5)P_2_ lipids fundamental to the functioning of eukaryotic membranes. Genetic perturbation that limit PI(4,5)P_2_ availability in early development are embryonic lethal, demonstrating the necessity of this lipid for living systems (13). Less severe mutations that modulate PI(4,5)P_2_ levels in cells underlie disease such as cardiomyopathies, cataract development, intellectual disability formation, and cancer (14, 15). Although the importance of PI(4,5)P_2_ lipids for cellular functions is self-evident, our understanding of how PI(4,5)P_2_ levels are stably maintained in cells is poorly understood.

Eukaryotes express two types of lipid kinases that regulate PI(4,5)P_2_ production: type I phosphatidylinositol-4-phosphate 5-kinase (PIP5K) enzymes and type II phosphatidylinositol-5-phosphate 4-kinase (PIP4K) enzymes. Type I PIP5K enzymes phosphorylate the 5-OH of a phosphatidylinositol-4-phosphate substrate (PI(4)P) (16, 17), while type II PIP4K enzymes utilize the less abundant phosphatidylinositol-5-phosphate (PI(5)P) lipid (18, 19). Despite the existence of these two pathways, studies have shown that PIP5K enzymes are primarily responsible for generating PI(4,5)P_2_ in mammalian cells (1, 20, 21). The PIP5K family consist of three paralogs (α, β, and γ), which exhibit tissue-specific expression patterns and serve non-redundant roles in cell signaling (21–23).

Cells lacking PIP4K paradoxically display increased PI(4,5)P_2_ levels (24). Recent biochemical studies revealed that the modulation of cellular PI(4,5)P_2_ levels depends on PIP4K-mediated inhibition PIP5K lipid kinase activity (25). Based on the ability of PIP5K to undergo membrane-mediated dimerization (26), we hypothesized that interactions between PIP4K or PIP5K paralogs modulate membrane-mediated dimerization of PIP5K to control PI(4,5)P_2_ lipid production. Supporting this model, cell biological studies have shown that overexpression of PIP5K, wild type or a catalytically dead mutant, increase cellular PI(4,5)P_2_ levels (25, 27). Under these conditions, overexpression of PIP5K is predicted to limit endogenous interaction with PIP4K, thus buffering the inhibitor nature of the PIP4K-PIP5K interaction. Although the interaction between PIP4K and PIP5K has been reported (24, 25, 28), the molecular basis remains unclear.

Membrane-mediated dimerization of PIP5K is transient and difficult to visualize by single molecule imaging. Quantifying the fraction of membrane dimerized PIP5K molecules in the presence of potential regulatory factors requires a new experimental approach. To address this challenge, we established a single molecule Forster resonance energy transfer (smFRET) assay to visualize homo- and heterodimerization of PIP5K paralogs on supported lipid bilayers by TIRF microscopy. Using this FRET sensor, we determined that membrane bound PIP4K interacts with the dimer interface of PIP5K, blocking membrane-mediated dimerization, which attenuates PIP5K lipid kinase activity. Guided by structural prediction, we generated PIP4K mutants that fail to block membrane-mediated dimerization or inhibit PIP5K lipid kinase activity in vitro. Expression of the PIP4K mutants in cells similarly fail to inhibit PIP5K-mediated production of the PI(4,5)P_2_ at the plasma membrane. Overall, this work reveals the molecular basis for PIP4K-mediated inhibition of PIP5K providing a new mechanism for understanding cellular PI(4,5)P_2_ lipid homeostasis.

## RESULTS

### Membrane-mediated dimerization of PIP5K visualized by smFRET

Membrane-mediated dimerization of PIP5K occurs in a protein density dependent manner on PI(4,5)P_2_ containing membranes (26). Membrane surface densities that favor PIP5K dimerization, however, are not well suited for direct visualization by dimerization using single molecule TIRF microscopy. As a result, we previously inferred PIP5K dimerization by monitoring changes in the dwell time and diffusion coefficient of Alexa Fluor 647 (AF647) PIP5K in the presence of unlabeled PIP5K (26).Alimitation of this approach is that interactions with other peripheral membrane binding proteins, such as PIP4K, can modulate the dwell time and diffusion coefficient of AF647-PIP5K in a manner that is indistinguishable from homodimerization of PIP5K. This presents a challenge interpreting whether a PIP4K regulates membrane-mediated dimerization of PIP5K or another biophysical property. Due to these limitations, we developed a single molecule FRET sensor to visualize membrane-mediated dimerization of PIP5K.

Homodimerization of PIP5KB is regulated by conserved residues (i.e. Asp-51 and Arg-254) that form salt-bridges at the dimer interface (**Figure 1A**). Using a combination of existing X-ray crystallography data (29, 30), AlphaFold structure prediction, and multiple sequence alignments of PIP5K family proteins, we identified candidate residues near the dimer interface that could potentially be labeled with fluorescent dyes to establish a FRET sensor for measuring dimerization (**Figure 1A**). Positioned in a solvent exposed loop near the dimer interface, we identified the non-conserved Pro-163 residue with a predicted separation of 11.5 Å at the PIP5KB dimer interface (**Figure 1A**). To engineer site-specific labeling, we identify all the solvent exposed cysteines through labeling of wild-type PIP5KB with Alexa Fluor 488 (AF488) maleimide. Electrospray ionization mass spectroscopy (ESI-MS) and accessible surface area calculations identified Cys-110 and Cys-411 as solvent exposed residues, while Cys-175 and Cys-256 were buried and protected from maleimide labeling. Based on these results, we mutated the solvent exposed residues (i.e. C110S, C411S) and introduced a P163C mutation for site-specific labeling near the dimer interface (**Figure 1A**). This mutant is referred to as the PIP5KB FRET construct. To confirm the mutations did not perturb the catalytic efficiency of PIP5KB, we performed lipid kinase assays to measure PI(4,5)P_2_ lipid production (**Figure 1B**). No significant change in the catalytic efficiency was observed for the PIP5KB FRET construct compared to the wild type kinase (**Figure 1C**).

**Figure 1.**
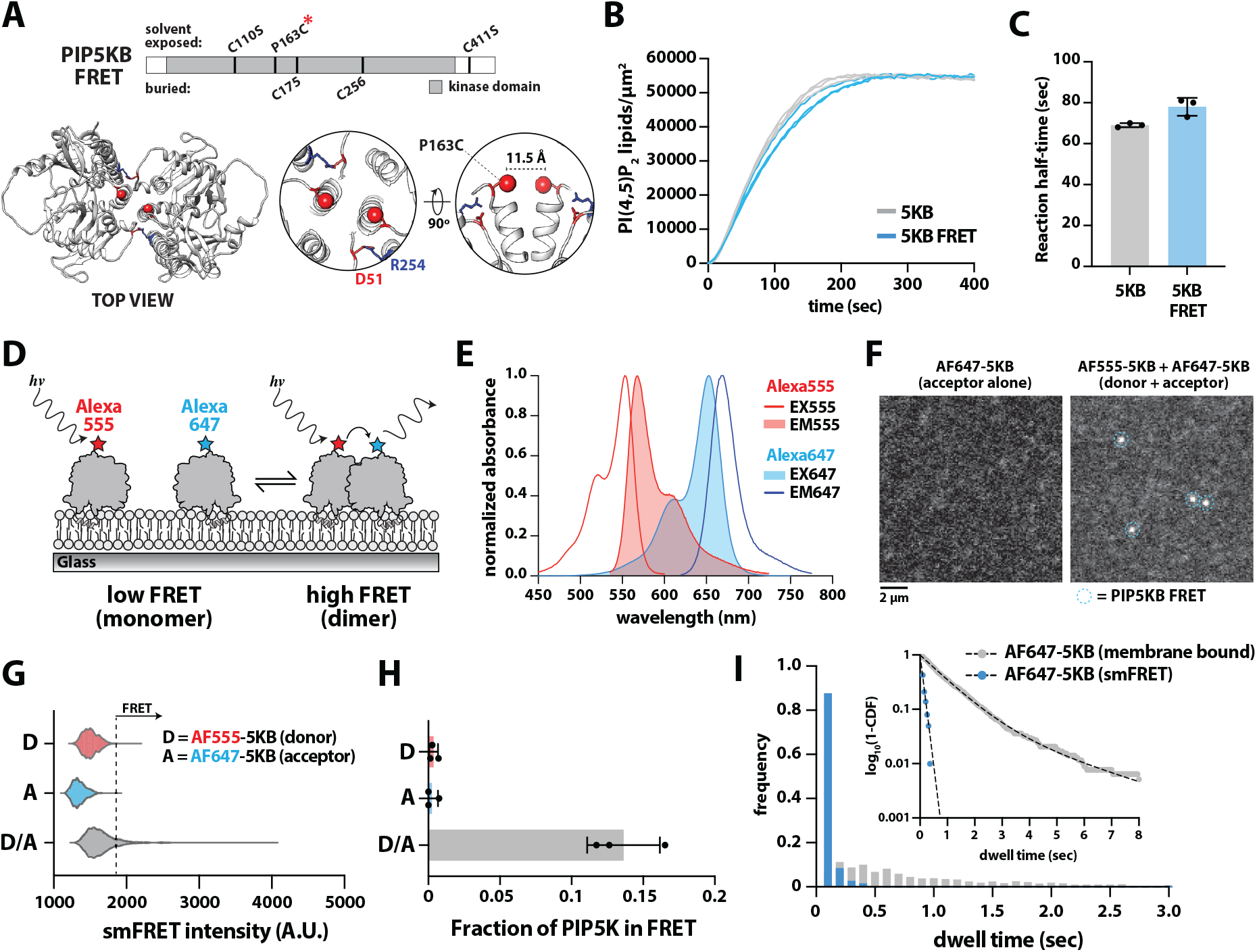
Membrane-mediated dimerization of PIP5K visualized by smFRET. **(A)** PIP5KB FRET construct indicated locations of buried (bottom) and solvent-exposed cysteines (top). AlphaFold structure prediction with the localization of dimer interface residues, D51 and R254. Red sphere indicates the location of P163C added for fluorophore labeling. Predicted distance between P163C residues in the dimeric is 11.5 Å. **(B)** Representative kinase activity trace showing the production of PI(4,5)P_2_ lipids in the presence of 20 nM Cy3-PLCδ PH and 5 nM PIP5KB, wild type and FRET construct mutant (C110S, P163C, C411S). **(C)** Quantification of reaction half-times comparing the activity of PIP5KB and PIP5KB FRET (n=3, p-value=0.41). **(D)** Cartoon schematic showing the molecular basis of smFRET based on membrane-mediated dimerization of PIP5KB. **(E)** Excitation-Emission spectra for AF555 and AF647 used for the FRET assay. **(F)** Representative TIRF-M images showing the sensitized acceptor emission of membrane bound AF647-PIP5KB in the presence of the AF555-PIP5KB acceptor. Data collected in the presence of either 1 nM acceptor alone (AF647-PIP5KB, left image) or 5 nM AF555-PIP5KB (donor) plus 1 nM AF647-PIP5KB (acceptor). **(G)** Molecular brightness distributions based on the fluorescence emission in the acceptor channel. Data collected in the presence 1 nM AF647-PIP5KB FRET acceptor alone, 5 nM AF555-PIP5KB FRET donor alone, or donor plus acceptor combined. Distributions contain the following number of data points: “D” (15 tracks, n = 39 spots); “A” (119 tracks, n = 622 spots); “D+A” (889 tracks, n = 2884 spots). These statistics were recorded over 60 seconds on a 3000 µm^2^ membrane surface. **(H)** Frequency of observing PIP5KB dimers by smFRET under the indicated conditions (*n*=3 replicates, bar equal SD). **(I)** Single molecule dwell time distribution for smFRET AF647-PIP5KB dimers (τ_1_=0.14 ± 0.03 sec, n=312 tracks, *N*=3 replicates) compared to the total membrane bound dwell time of AF647-PIP5KB (τ_1_=0.89 ± 0.03 sec, τ_2_=2.7 ± 0.3 sec, α=0.94, n=2362 tracks, *N*=3 replicates). Alpha(α) represents the fraction of molecules with the time constant (τ_1_). Plot inset is log_10_(1-cumulative distribution frequency) with fit (dashed line) to single or double exponential. Membrane compositions: **(B-C)** 2% PI(4) P, 98% DOPC; **(F-I)** 4% PI(4,5)P_2_, 96% DOPC.

To establish the smFRET assay to visualize membrane-mediated dimerization, we labeled purified PIP5KB FRET construct (referred to as PIP5KB) with either Alexa Flour 555 (AF555) or Alexa Fluor 647 (AF647) (**Figure 1D**). These fluorescent dyes have high quantum yields, robust photostability, and a FÖrster Radius (R_0_) of 51 Å, making them an ideal pair for smFRET (**Figure 1E**). Using Total Internal Reflection Fluorescence Microscopy (TIRF-M), we visualized the sensitized emission of AF647-PIP5KB (acceptor) with single molecule resolution in the absence and presence AF555-PIP5KB (donor) on supported lipid bilayers (SLBs) containing PI(4,5)P_2_ lipids (**Figure 1F**). Conditions were optimized to excite AF555-PIP5KB and visualize AF647-PIP5KB emission with minimal direct excitation of AF647 using either a 532 or 561 nm laser (**Figure S1**). Under these conditions, a distinct population of bright AF647-PIP5KB molecules were observed only in the presence of both the donor and acceptor labeled PIP5KB (**Figure 1F-1G** and **Movie S1**). The frequency of observing PIP5KB dimers by smFRET was calculated based on a threshold molecular brightness of visualized AF647-PIP5KB (acceptor) that exceeded the brightest particles in our control conditions (**Figure 1H**). Comparing the FRET lifetime to the total membrane dwell times of AF647-PIP5KB, revealed that the lifetime of PIP5K homodimers was relatively transient (τ_1_=0.14 ± 0.03 sec) compared to the total membrane dwell time of a single membrane bound kinase (τ_1_=0.89 ± 0.03 sec, τ_2_=2.7 ± 0.3 sec, α=0.94) (**Figure 1I**).

### Mutational analysis and competition validate specificity of smFRET assay

Membrane-mediated dimerization of PIP5K is regulated by salt bridges formed between aspartic acid and arginine residues (**Figure 1A**). Two previous studies showed that mutations in these residues disrupt homodimerization and attenuate PIP5K lipid kinase activity (26, 29). To validate the specificity of the smFRET PIP5KB dimerization sensor, we characterized a dimerization deficient PIP5KB mutant, D51R (**Figure 2A**). The D51R mutation was introduced into the PIP5KB FRET construct and labeled with the acceptor fluorophore (AF647). We performed the smFRET assay with a wild type AF555-PIP5KB (donor) and mutant AF647-PIP5KB (D51R, acceptor).

**Figure 2.**
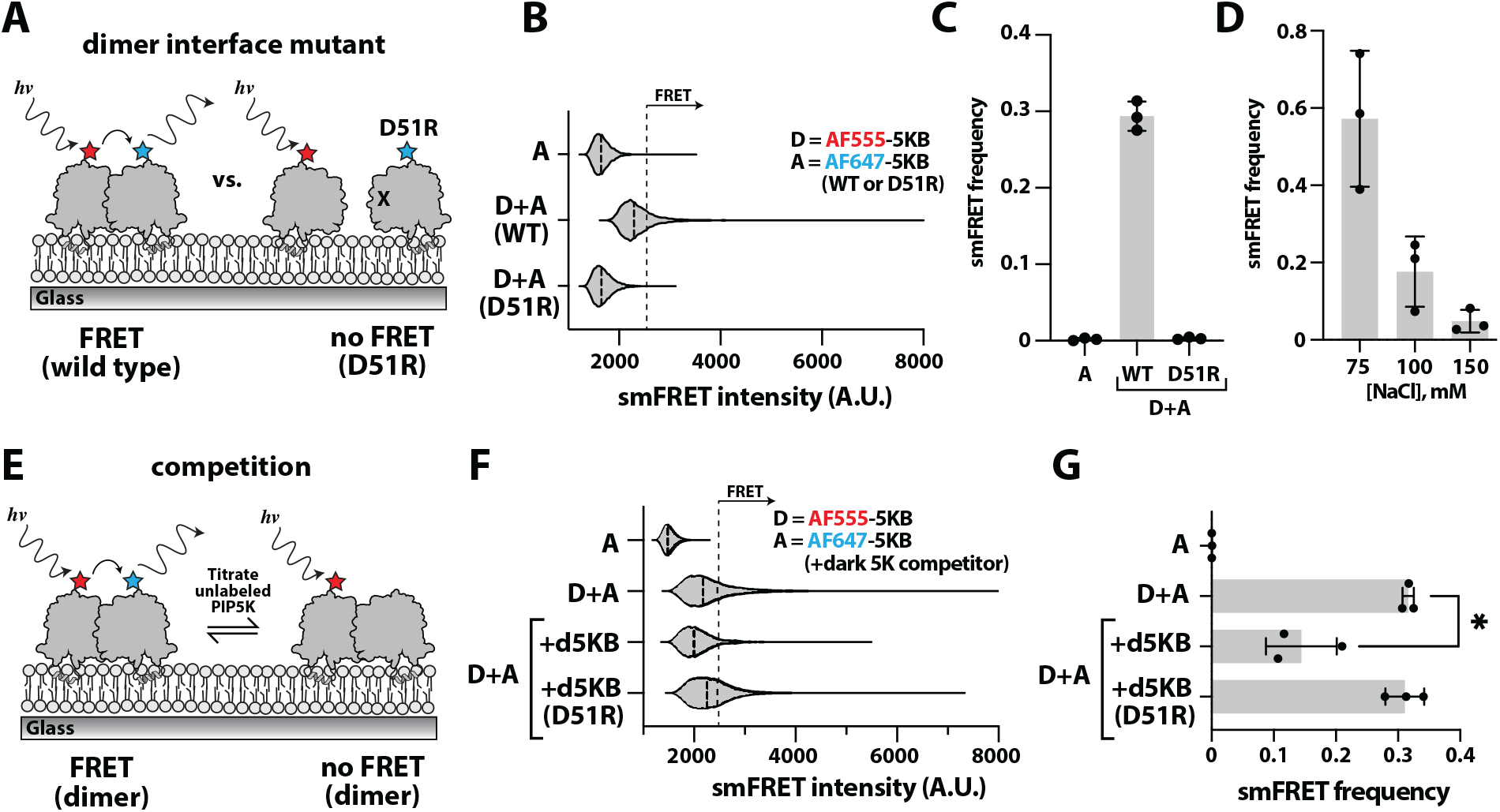
Dimer interface mutation and competition reduce PIP5KB smFRET efficiency. **(A)** Cartoon schematic showing lack of smFRET measured in the presence of AF555-PIP5KB (WT) and AF647-PIP5KB (D51R) dimer interface mutant. **(B)** Representative smFRET particle intensity distributions in the presence of 5 nM AF555-PIP5KB and 1 nM AF647-PIP5KB (WT or D51R). **(C)** Quantification of frequency particles exhibit smFRET in (B). *p=0*.*0008* comparing “WT” and “D51R” conditions. **(D)** Lower ionic strength buffer increases smFRET efficiency. Data collect using 5 nM AF555-PIP5KB plus 1 nM AF647-PIP5KB in the presence of varying NaCl concentrations (*p=0*.*034* comparing 75 mM and 150 mM conditions). **(E)** Cartoon schematic showing how competition with unlabeled PIP5K is predicted to decrease the smFRET efficiency between AF555-PIP5KB (donor) and AF647-PIP5KB (acceptor). **(F)** Representative smFRET particle intensity distributions measured in the absence and presence of 10 nM unlabeled PIP5KB competitor (WT or D51R). **(G)** Quantification of frequency that particles exhibit smFRET in (F). n=3 technical replicates for each experiment shown. Bars equal SD. Membrane composition: 4% PI(4,5)P_2_, 96% DOPC.

Under these conditions, we observed a significant decrease in the frequency of smFRET compared to the wild type PIP5KB FRET construct (**Figure 2B-2C**). Consistent with PIP5K dimerization requiring electrostatic interactions between kinase domains, the frequency of observing membrane bound PIP5KB dimers by smFRET was inversely proportional to the buffer ionic strength (**Figure 2D**). PIP5KB dimers detected by smFRET also exhibited a characteristic decrease in membrane diffusivity compared to the monomeric species (**Figure S2A-B**). This change in membrane binding behavior occurs in PIP5K density-dependent manner and requires an intact dimer interface (26). Overall, these results validate our smFRET approach enabling the quantification of PIP5K dimerization across experimental conditions to determine how various protein-protein interactions modulate membrane-mediated homodimerization of PIP5K.

In vivo, PIP5K associates with the plasma membrane where competition for protein-protein and protein-lipid interactions is predicted to modulate membrane-mediated dimerization of PIP5K. In our smFRET assay, competition with unlabeled PIP5KB is expected to reduce the probability of dimerization between donor and acceptor labeled PIP5KB if membrane-bound dimers are in dynamic equilibrium with monomeric species (**Figure 2E**). Consistent with this prediction, the addition of unlabeled PIP5KB significantly reduced the frequency of particles observed by FRET, while the addition of the PIP5KB (D51R) dimer interface mutant was unable to promote FRET (**Figure 2F-2G**). In contrast, experiments performed in the presence of a peripheral membrane binding protein with broad specificity to anionic lipids in vitro (i.e. LactC2) was unable to disrupt membrane-mediated dimerization of PIP5KB observed by smFRET (**Figure S3**). Together, these results demonstrate that an intact dimer interface is required to observe PIP5KB smFRET in vitro. In addition, AF555-PIP5KB does not support FRET with membrane bound AF647-LactC2.

Overexpression of catalytically dead PIP5K paradoxically increases the total plasma membrane density of PI(4,5)P_2_ lipids in cells (25, 27). One hypothesis is that catalytically dead PIP5K promotes membrane-mediated dimerization of endogenously expressed PIP5K. To test this hypothesis, we performed smFRET using donor labeled AF555-PIP5KB FRET (D266K). We previously showed that the D266K mutant binds to PI(4,5)P_2_ containing membranes similar to the wild type kinase in vitro (31). Using AF555-PIP5KB (D266K) as the FRET donor, we detected a similar frequency of smFRET events compared to the wild type PIP5KB donor and acceptor pair (**Figure S4**). These data suggest that overexpression of catalytically dead PIP5KB promotes membrane-mediated dimerization of endogenously expressed PIP5K, which stimulates lipid kinase activity and increases cellular PI(4,5)P_2_ levels.

### Membrane-mediated heterodimerization of PIP5K paralogs

PIP5K paralogs share a conserved dimer interface that is expected to facilitate membrane-mediated heterodimerization (**Figure 3A**). Since PIP5K paralogs have similar tissue expression profiles (32), heterodimerization of paralogs could provide a mechanism that modulates cellular PI(4,5)P_2_ levels. Previous efforts to detect heterodimerization of PIP5K paralogs have relied on co-immunoprecipitation (33) and colocalization visualized at the cellular plasma membrane (3, 25, 34). Here, we tested whether purified PIP5KA and PIP5KB can heterodimerize on SLBs using single molecule TIRF-M. Previously, we established that dimeric PIP5KB has distinct single molecule membrane binding properties compared to monomeric PIP5KB (26). By comparing the membrane dwell time and diffusion coefficient of AF647-PIP5KB in the absence and presence of dark unlabeled PIP5K (**Figure 3A**), we detected membrane-mediated homodimerization of PIP5KB (26). Repeating these experiments in the presence of PIP5KA, revealed a similar increase in membrane dwell time and reduction in membrane diffusion for AF647-PIP5KB (**Figure 3B-3C**). We previously showed that these changes in membrane dwell time and lateral diffusion are not observed for fluorescently labeled PIP5K containing a dimer interface mutation (26). Together, these observations indicate that PIP5K paralogs can heterodimerize on membranes.

**Figure 3.**
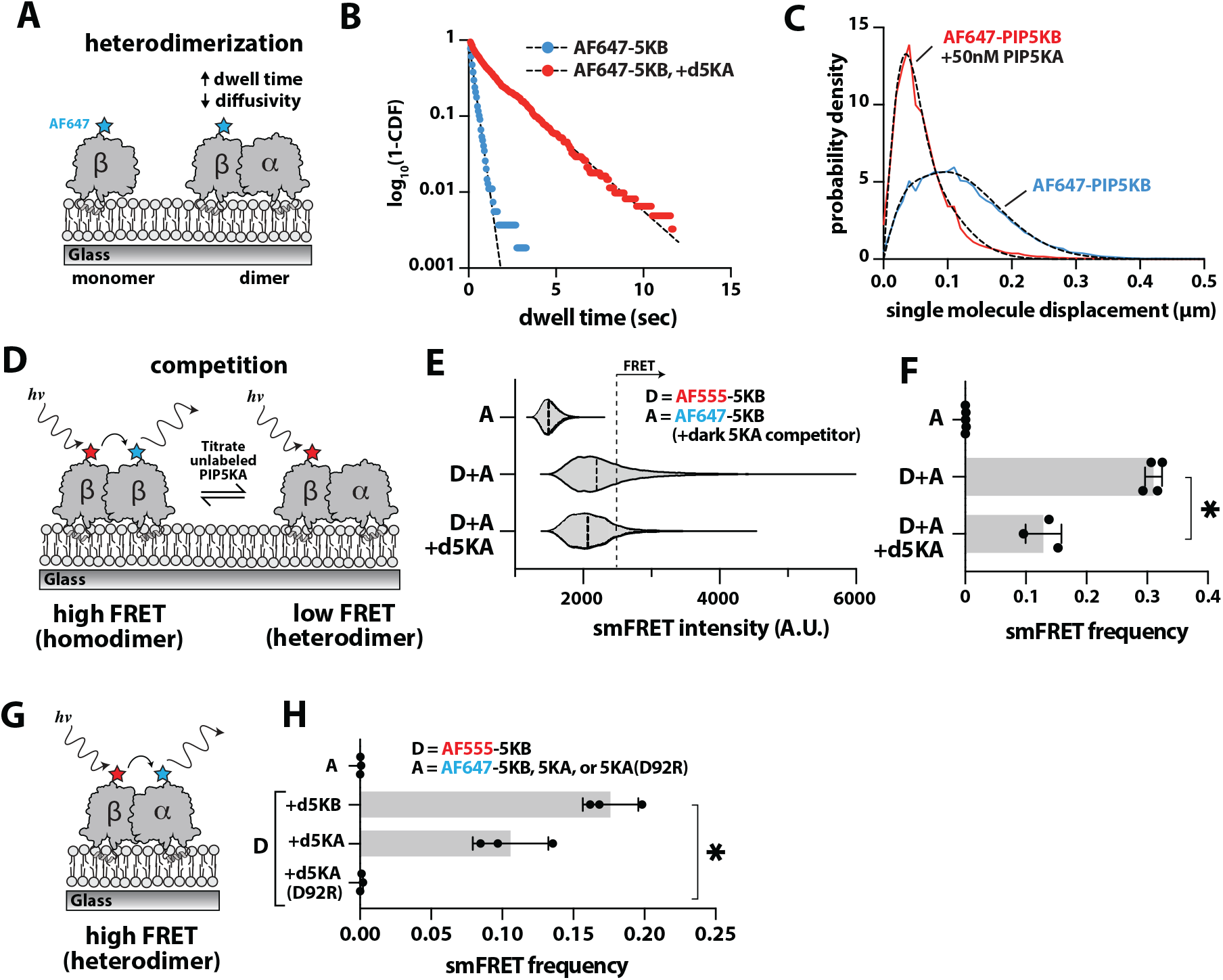
Membrane-mediated heterodimerization of PIP5K paralogs. **(A)** Cartoon schematic showing how heterodimerization is predicted to increase the membrane dwell time and decrease diffusion. **(B)** PIP5KA increases the single molecule dwell time of AF647-PIP5KB. Representative log_10_(1-cumulative distribution frequency) dwell time distributions measured in the presence of 10 pM AF647-PIP5KB alone (blue; τ_1_=0.28 ± 0.02 sec, N=1483 tracks, *n*=3 replicates) or in the presence of 50 nM unlabeled PIP5KA (red; τ_1_=0.59 ± 0.14 sec, τ_2_=2.19 ± 0.36 sec, α=0.45, N=1791 tracks, *n*=3 replicates). **(C)** Heterodimerization with PIP5KA reduces the diffusivity of membrane bound AF647-PIP5KB. Representative step size distributions measured in the presence of 10 pM AF647-PIP5KB alone (blue) or in the presence of 50 nM unlabeled PIP5KA (red). **(D)** Schematic showing that heterodimerization between AF647-PIP5KB and unlabeled PIP5KA is predicted to eliminate smFRET signal. **(E)** Representative maximum particle intensities distributions measured in the presence of 5 nM AF555-PIP5KB and 1 nM AF647-PIP5KB (“D+A”) or plus the addition of 10 nM unlabeled PIP5KA (“D+A+d5KA”). **(F)** Quantification of frequency particles exhibit smFRET in (E). Bars equal SD, n=3-5 technical replicates (\**p=0*.*0036*). **(G)** Cartoon schematic showing smFRET based on heterodimerization between AF555-PIP5KB and AF647-PIP5KA. (H) Quantification of smFRET frequency measuring between the indicate proteins. AF647-PIP5KA dimer interface mutant (D29R) does not exhibit smFRET based on heterodimerization with AF555-PIP5KB, while wild type AF647-PIP5KA does. Donor and acceptor concentration equal 5 nM and 1 nM, respectively. Bars equal SD, n=3 technical replicates (\**p=0*.*021*). Membrane composition: 4% PI(4,5)P_2_, 96% DOPC.

Having established that the smFRET assay detects homodimerization of PIP5KB, we next tested whether competition via heterodimerization with PIP5KA could reduce the frequency of PIP5KB homodimerization (**Figure 3D**). Like the competition experiments that utilized dark unlabeled PIP5KB (**Figure 2E-2G**), the addition of unlabeled PIP5KA blocked homodimerization of the PIP5KB FRET construct on SLBs (**Figure 3E-3F**). To determine whether we could directly visualize heterodimerization by smFRET, we combined AF555-PIP5KB (donor) and AF647-PIP5KA (acceptor) on PI(4,5)P_2_ containing membranes (**Figure 3G**). Like the PIP5KB homodimerization FRET sensor, we visualized a high frequency of PIP5KA-PIP5KB heterodimers by smFRET compared to the acceptor alone control (**Figure 3H**). To validate the specificity of the heterodimer FRET signaling, we repeated the experiments using the dimerization deficient mutant AF647-PIP5KA (D92R). Similar to measurements performed with AF647-PIP5KB (D51R) (**Figure 2A-2C**), smFRET based on heterodimerization was rarely detected in the presence of the AF647-PIP5KA (D92R) mutant acceptor (**Figure 3H**).

### PIP4K attenuates PIP5K activity by blocking membrane-mediated dimerization

Type II phosphatidylinositol-5-phosphate 4-kinase (PIP4K) family produce PI(4,5)P_2_ by phosphorylating PI(5)P lipids (35). Due to the low abundance of PI(5)P in cells, however, this pathway contributes only a minor fraction of the total plasma membrane pool of PI(4,5)P_2_. Paradoxically, genetic knock out all three PIP4K paralogs in human cells cause cellular PI(4,5)P_2_ levels to significantly increase (24). This affect has been attributed to PIP4K’s ability to directly bind PIP5K and attenuate lipid kinase activity (24, 25). The mechanism of PIP4K-mediated inhibition remains unclear.

Due to differences in their specificity loop sequences, purified PIP4K and PIP5K require different densities of PI(4,5)P_2_ lipids for membrane localization in vitro (25, 31). Leveraging this difference in membrane avidity, we quantified the ability of PIP5KB to directly bind and recruit AF488-PIP4KA to membranes containing 2% PI(4,5)P_2_ (**Figure 4A**). Under these conditions, AF488-PIP4KA alone remains primarily in the solution phase with minimal membrane localization (**Figure 4B-4C**). Following the addition of PIP5KA, however, we observed robust membrane recruitment of AF488-PIP4KA (**Figure 4B-4C**). In contrast, spike in experiments performed with the dimerization deficient PIP5KA (D92R) mutant failed to enhance AF488-PIP4KA membrane localization (**Figure 4B-4D**). We observed similar results following the addition of PIP5KB, wild type or a dimerization deficient mutant (i.e. D51R) (**Figure 4D**).

**Figure 4.**
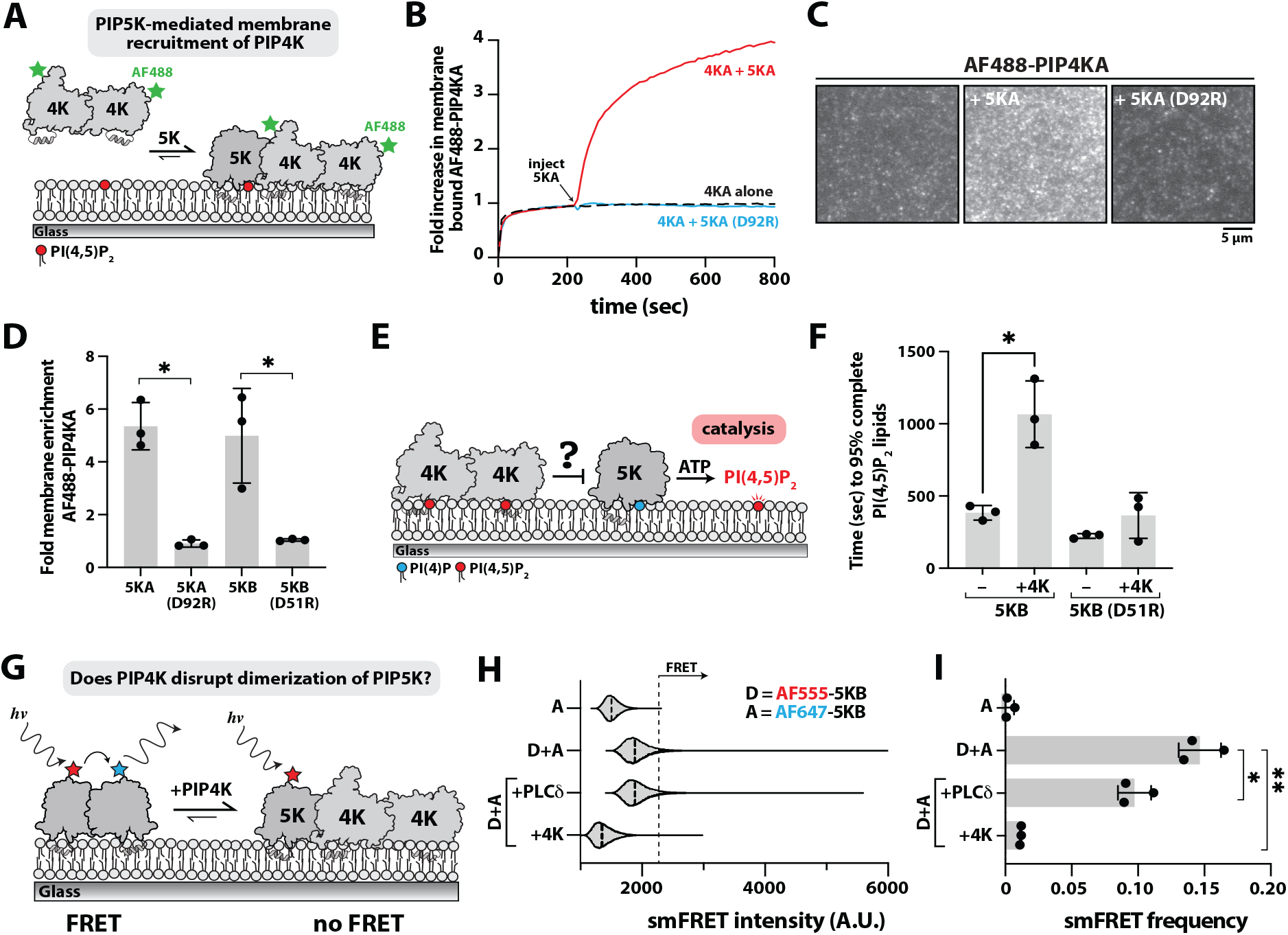
PIP4K disrupts homodimerization of PIP5K. **(A)** Cartoon illustrating showing that PIP4K localization to membranes containing 1% PI(4,5)P_2_ requires PIP5K. **(B)** Kinetic traces showing membrane recruitment of 50 nM AF488-PIP4K following the addition of 10 nM PIP5KA. PIP5KA-mediated membrane recruitment of AF488-PIP4K is not observed in the presence of 30 nM PIP5KA (D92R) dimer interface mutant. **(C)** Representative TIRF-M images showing plateau equilibrium membrane localization of 50 nM 488-PIP4KA in the absence or presence of either 10 nM PIP5KA or 30nM monomeric PIP5KA (D92R). **(D)** Quantification of fold change in 50 nM AF488-PIP4KA membrane recruitment in the presence of 10nM PIP5KA, 30nM PIP5KA D92R (\**p= 0*.*007*), 10nM PIP5KB, or 30nM PIP5KB D51R (\**p=0*.*009*). Bars equal SD, n=3 technical replicates. Membrane composition: 2% PI(4,5)P_2_, 98% DOPC. **(E)** Cartoon illustrating SLB assay for measuring PIP4K-mediated inhibition of PIP5K lipid kinase activity. **(F)** PIP4K attenuates PIP5KB activity. Quantification of time required for PIP5K to generate 95% of maximum PI(4,5)P_2_ lipid density. Kinase activity measured in the presence of 1 nM PIP5KB +/− 50 nM PIP4KA (\**p=0*.*031*) or 10 nM PIP5KB (D51R) +/− 50 nM PIP4KA (*p=0*.*26*). Reaction monitored in the presence of 20 nM Cy3-PLCδ PH domain. n=3 technical replicates. Bars equal SD. Membrane composition: 4% PI(4)P, 96% DOPC. **(G)** Experimental design for determining whether PIP4K blocks membrane-mediated dimerization PIP5K based on smFRET. **(H)** Representative smFRET intensities distributions measured in the presence of 5 nM AF555-PIP5KB (donor) and 1 nM AF647-PIP5KB (acceptor). Competition for PIP5K homodimerization was measured in the presence of 50 nM PIP4KA or 50 nM PLCδ PH domain. **(I)** Quantification of smFRET efficiency measured in (H). PIP4K reduces the smFRET efficiency of PIP5K (\*\**p=0*.*014*), but not the PLCδ PH domain (\**p=0*.*1*). Bars equal SD, n=3 technical replicates. Membrane composition: 4% PI(4,5)P_2_, 96% DOPC.

Since membrane-mediated dimerization enhances PIP5K catalytic activity (26), we hypothesized that PIP4K blocks homodimerization and attenuates PIP5K activity (**Figure 4E**). If PIP4K disrupts membrane-mediated dimerization, then a dimerization deficient PIP5K mutant should be insensitive to PIP4K inhibition of kinase activity. To test this hypothesis, we measured PIP5K activity in the absence and presence of a physiologically relevant concentration of PIP4KA (**Figure 4E-4F**). As previously reported (25), the presence of PIP4KA increased the time required for PIP5KB to generate PI(4,5)P_2_ on SLBs by ~3-fold (**Figure 4F**). In contrast, the PIP4K-dependent decrease in the rate of PI(4,5)P_2_ production was minor in the presence of monomeric PIP5KB (D51R) (**Figure 4F**). This small decrease is likely due to PIP4KA binding to PI(4,5)P_2_ lipids and slightly reducing membrane localization of PIP5KB (D51R) as more PI(4,5)P_2_ is generated. Together, these results suggest that PIP4K binds to the PIP5K dimer interface, thus creating a PIP5K-(PIP4K)_2_ (PIP4K is a constitutive dimer) ternary complex that is less catalytically activity compared to homodimeric PIP5K.

To test whether PIP4K blocks membrane-mediated dimerization of PIP5K, we performed smFRET competition experiments (**Figure 4G**). Consistent with our previous results, homodimerization of PIP5KB measured by smFRET showed a significant decrease in the FRET frequency in the presence of PIP4KA (**Figure 4H-4I**). While 50 nM PIP4KA is a physiologically relevant concentration of protein in cells, it’s possible that the decrease in FRET we see is due to molecular crowding on our supported bilayer, rather than a direct PIP4K-PIP5K interaction. As a control, we performed the smFRET assay in the presence of the PI(4,5)P_2_ lipid binding domain, PLCδ PH domain, which bind membranes but does not associate with either the acceptor or donor labeled PIP5KB. We found that PLCδ PH resulted in a small decrease in the smFRET efficiency (**Figure 4H-4I**). Together, this shows that the decrease we see from the addition of PIP4KA in our FRET assay results from direct blockage of the dimer interface and not solely based on competition for PI(4,5)P_2_ lipid binding sites. Taken together, these data support the hypothesis that PIP4K inhibits PIP5K activity by blocking PIP5K dimerization by competing for binding at the dimer interface.

### Molecular basis of PIP4K-PIP5K interaction

To map the interaction site between PIP4K and PIP5K, we modeled potential binding sites using AlphaFold multimer (36) (**Figure 5A**). Based on this structure prediction, M60/L61/M62 residues in PIP4KA were predicted to form a hydrophobic interaction at the dimer interface of PIP5K (**Figure 5A** and **Figure S5**). To determine whether these residues are required for PIP4K-mediated inhibition of PIP5K, we measured kinase activity on supported membranes. Consistent with our previous observation (25), the presence of wild type PIP4K attenuated PIP5KB activity after a threshold density of ~3% PI(4,5)P_2_ was generated (**Figure 5B**). By contrast, the PIP4KA (M60R, L61E, M62R) mutant was unable to inhibit PIP5K lipid kinase activity (**Figure 5B-5C**). The AlphaFold multimer also predicted an interaction between PIP5K with PIP4KA R104 (**Figure S6A**). However, the PIP4KA (R104D) mutant was still able to interact with PIP5K and inhibit lipid kinase activity like wild type PIP4K (**Figure S6**).

**Figure 5.**
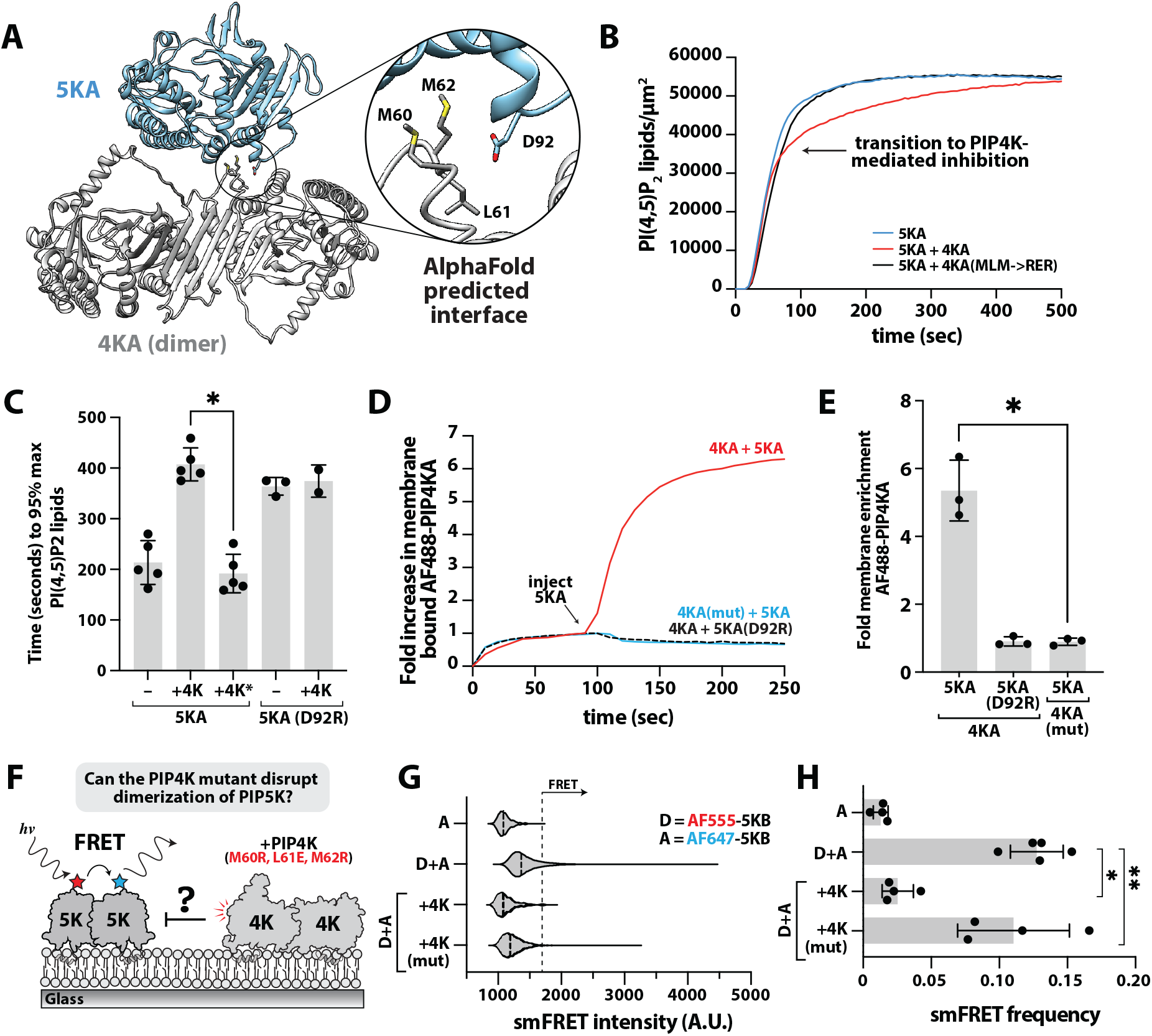
Molecular basis of PIP4K-PIP5K interaction. **(A)** PIP4K-PIP5K binding interaction based on Alphafold3 structure prediction. **(B)** Representative kinase activity trace monitoring the production of PI(4,5)P_2_ lipids in the presence of 20 nM Cy3-PLCδ, 1 nM PIP5KA, and 50 nM PIP4KA (wild type or M60R, L61E, M62R mutant). **(C)** Quantification of time required for PIP5K to generate 95% of maximum PI(4,5)P_2_ lipid density in the presence of indicated proteins. All experiments include 1 nM PIP5KA (wild type or D92R) +/− 50 nM PIP4K (wild type or M60R, L61E, M62R mutant). Bars equal SD, n=2-5 technical replicates (\**p<0*.*0001*). Membrane composition: 4% PI(4,5)P_2_, 96% DOPC. **(D)** PIP5KA is unable to promote membrane recruitment of AF488-PIP4KA (M60R, L61E, M62R). Enhanced membrane recruitment of 50 nM AF488-PIP4KA (wild type and mutant) was monitored following the injection of 10 nM PIP5KA (wild type or D92R). **(E)** Quantification of fold change in AF488-PIP4KA membrane recruitment following the addition of PIP5KA in (D). Bars equal SD, n=3 technical replicates (\**p=0*.*012*). Membrane composition: 2% PI(4,5)P_2_, 98% DOPC. **(F)** Experimental design for measuring whether PIP4K (M60R, L61E, M62R) inhibits membrane-mediated dimerization of PIP5K measured by smFRET. **(G)** Representative smFRET intensities distributions measured in the presence of 5 nM AF555-PIP5KB (donor) and 1 nM AF647-PIP5KB (acceptor). Competition for PIP5K homodimerization based on smFRET measured in the presence of 50 nM PIP4KA (wild type or M60R, L61E, M62R). **(H)** Quantification of smFRET efficiency measured in (G). PIP4K reduces the smFRET efficiency of PIP5K (\**p<0*.*0001*), but not PIP4KA (M60R, L61E, M62R) (\*\**p=0*.*487*). Bars equal SD, n=4-5 technical replicates. Membrane composition: 4% PI(4,5)P_2_, 96% DOPC.

To determine how mutations in PIP4K modulate interactions with PIP5K, we quantified the ability of membrane bound PIP5K to recruit AF488-PIP4K to membranes containing 1% PI(4,5)P_2_. Consistent with the mutant PIP4KA being unable to interact with PIP5KA, we did not observe PIP5K-mediated membrane recruitment of AF488-PIP4KA (M60R, L61E, M62R) (**Figure 5D-5E**). This result mimics the inability of dimerization deficient PIP5K (D92R) to enhance membrane recruitment of wild type AF488-PIP4K (**Figure 5D-5E**). Using the smFRET assay, we tested whether PIP4KA (M60R, L61E, M62R) could block membrane-mediated dimerization of PIP5KB (**Figure 5F**). Consistent with our previous results, the frequency of PIP5KB homodimerization was significantly reduced in the presence of PIP4KA (**Figure 5G-5H**). However, the presence of PIP4KA (M60R, L61E, M62R) did not significantly change the frequency of PIP5KB smFRET (**Figure 5H**). To ensure that differences observed between wild-type and mutant PIP4KA could not be attributed protein misfolding, we measured the secondary structure content of PIP4K using circular dichroism (CD). This yielded CD spectra that were indistinguishable comparing PIP4KA (M60R, L61E, M62R) to the wild type kinase (**Figure S7A-S7B**). The size exclusion chromatography elution profiles comparing wild type and mutant PIP4K were also nearly identical (**Figure S7C**). To determine whether the PIP4KA (M60R, L61E, M62R) displayed similar membrane binding properties, we measured cooperative membrane binding to PI(4,5)P_2_ lipids generated by the yeast PIP5K homolog, Mss4 (**Figure S9A**). As previously reported (25), both AF488-labeled PIP4K proteins exhibited half maximal membrane binding when the molar density of PI(4,5)P_2_ reached ~3% (**Figure S9B-S9C**).

### Disruption of the PIP4K-PIP5K interface prevents PIP4K modulation of PI(4,5)P_2_ in cells

Over-expression of any PIP5K paralog in cells increases the density of PI(4,5)P2 at the plasma membrane, whereas co-overexpression of any PIP4K paralog suppresses this effect (25). We therefore tested if the PIP4KA-PIP5KA interface that we identified here was necessary to suppress PIP5K-mediated production of PI(4,5)P_2_ at the plasma membrane (PM). As expected, over-expression of GFP-PIP5KA increased PI(4,5)P_2_ as measured with the low affinity biosensor, Tubby_c_^R332H^ (37). This effect was reversed by co-overexpression of BFP-PIP4KA (**Figure 6A-6B**). By contract, a mutated BFP-PIP4KA (M60R,L61E,M62R; denoted 4K*) failed to attenuate the PIP5KA-mediated increase in PI(4,5)P2 levels (**Figure 6A-6B**). We made an additional PIP4KA mutant denoted 4KΔ, whereby the hydrophobic loop containing M61-M62 was truncated, replacing residues 54-62 with a short GGSGG linker. This construct also failed to reverse the PIP5KA-mediated increase in PM PI(4,5)P_2_ (**Figure 6A-6B**). To study the effects of expressed PIP4Ks on endogenous PIP5Ks, we previously generated a myristoylated PIP4KA for constitutive plasma membrane targeting (25). This construct reduces steady-state PI(4,5)P2 in cells. We once again observed a reduction in PM PI(4,5)P_2_ levels in cells expressing myristoylated PIP4KA, but not when we introduced the 4K* mutation or 4KΔ truncation (**Figure 6C-6D**).

**Figure 6.**
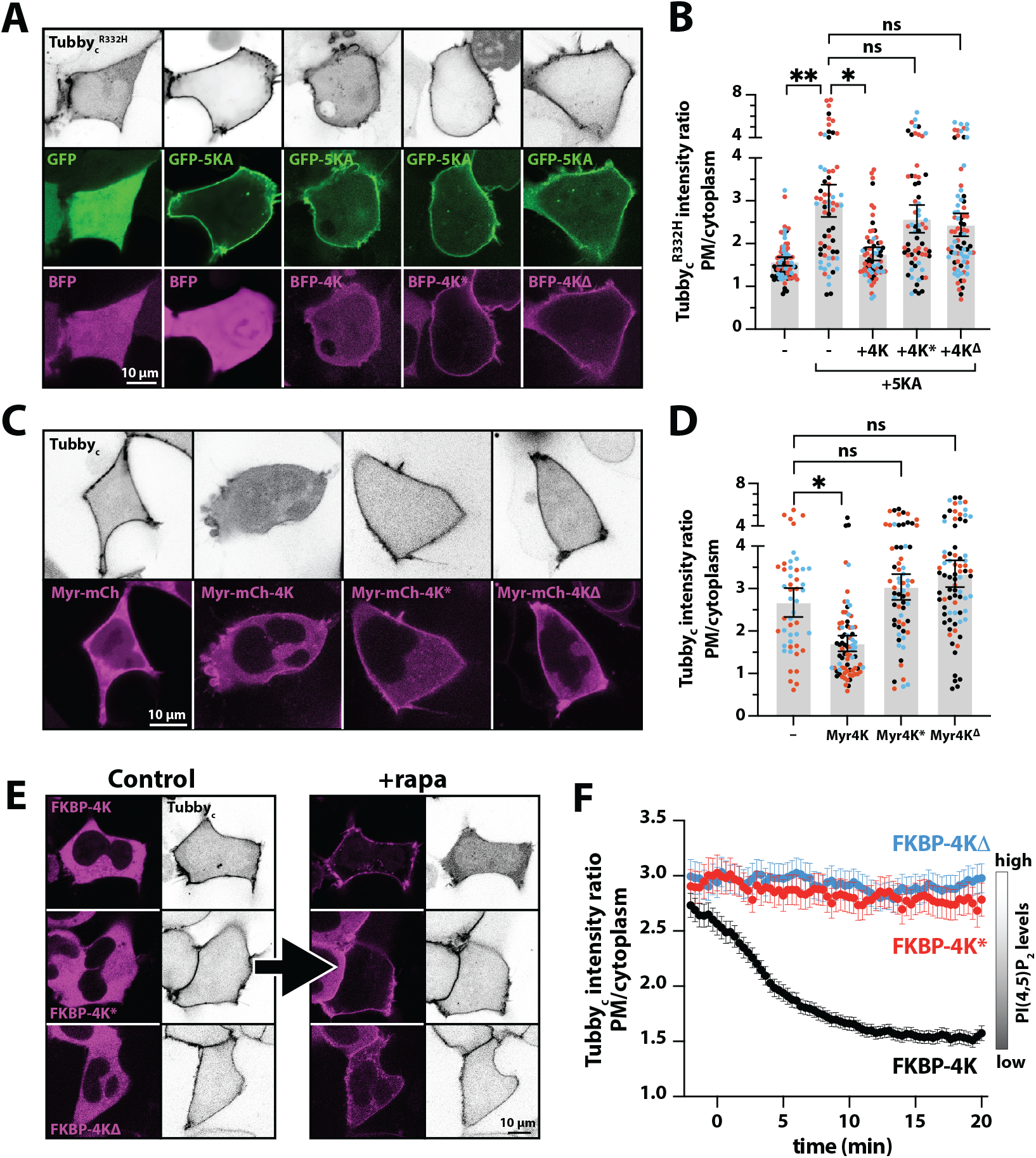
PIP4K-PIP5K interaction is required to modulate PI(4,5)P_2_ synthesis in cells. **(A)** Representative confocal sections of HEK293A cells expressing low affinity PI(4,5)P_2_ biosensor Tubby _c_^R332H^-mCherry and either GFP or GFP-PIP5KA plus BFP, BFP-PIP4KA (4K), BFP-PIP4KA^M60R, L61E, M62R^ (4K*) or BFP-PIP4KA-Δ54-62, where residues H54-M62 are replaced with a short GGSGG linker (4KΔ). **(B)** Quantification of the fluorescence ratio of Tubby _c_^R332H^-mCherry at the plasma membrane relative to cytoplasm from the same experiment as (A). Bars are means ± 95% C.I. of all cells acquired from three experiments, whereas individual data points show individual cell data color coded by experimental replicate. Statistics show the results of a Šídák’s multiple comparisons test after nested 1-way ANOVA (*p=0*.*0054*l *F=7*.*197, DF*_*n*_*=4; Df*_*d*_*=10; three experimental replicates with 15-33 cells per replicate)*. \*\**p=0*.*0048*; \**p=0*.*0118*. **(C)** Representative confocal sections of HEK293A cells expressing high affinity PI(4,5)P_2_ biosensor Tubby_c_-mCherry and either myristolated mCherry (Myr-mCh), Myr-mCh-4KA, Mry-mCh-4KA* or Myr-4KΔ. **(D)** Quantification of the fluorescence ratio of Tubby_c_-mCherry at the plasma membrane relative to cytoplasm from the same experiment as (C). Bars are means ± 95% C.I. of all cells acquired from two or three experiments, whereas individual data points show individual cell data color coded by experimental replicate. Statistics show the results of a Šídák’s multiple comparisons test after nested 1-way ANOVA (*p=0*.*0017*l *F=15*.*71, DF*_*n*_*=3; Df*_*d*_*=7; three experimental replicates with 17-39 cells per replicate)*. \**p=0*.*0308*. **(E)** Representative confocal sections of HEK293A cells expressing high affinity PI(4,5)P_2_ biosensor Tubby_c_-mCherry, Lyn_11_-FRB-iRFP (not shown) and either mCherry-tagged FKBP-4K, -4K* or 4KΔ. Images are shown before or 20 min after the addition of 1 µM rapamycin to induce FRB-FKBP heterodimerization and PM recruitment of the 4K constructs. **(F)** Quantification of time lapse experiments for the experiment from (E), showing the fluorescence ratio at the plasma membrane relative to cytoplasm for Tubby_c_. Data are means ± s.e.m. of 36-78 cells from 3-7 experiments.

A potential trivial explanation for the failure of mutant PIP4KA to suppress PIP5K in cells is that the proteins adopt a non-functional conformation. To rule this out, we capitalized on the selectivity loop mutant A371E in PIP4KA, which was shown to swap enzyme activity from a PI(5)P 4-kinase to become a PI(4)P 5-kinase (38). A fusion of this mutant PIP4KA to FKBP (FKBP-4K^5K^) could be recruited to the plasma membrane by rapamycin-induced dimerization with a co-expressed FRB domain, targeted by a palmitoylated and myristoylated N-terminal peptide derived from Lyn kinase (**Figure S9B-S9C**). This recruitment led to a robust increase in plasma membrane PI(4,5)P_2_ detected with low affinity Tubby_c_^R332H^ biosensor (**Figure S9D**). We next introduced the 4K* mutations or 4KΔ truncation into this A371E mutant. The mutant kinases recruited as efficiently as the unmutated enzyme (**Figure S9C**) and retained robust PI(4)P 5-kinase activity (**Figure S9D**). Consistent with our in vitro measurements (**Figure S7-S8**), the PIP5K interaction-disrupting mutations still allowed PIP4K to adopt a functional kinase conformation in vivo. Next, we introduced the 4K* and 4KΔ mutations into the wild-type PI(5)P 4-kinase background. Whereas acute PM recruitment of FKBP-PIP4KA showed a potent reduction of PI(4,5)P_2_ in HEK293A cells (**Figure 6E-6F**), as we have previously shown (25), neither mutant was able to reduce PI(4,5)P_2_ levels (**Figure 6E-6F**) despite equally robust plasma membrane recruitment (**Figure S9E**).

## DISCUSSION

We developed a novel smFRET sensor for visualizing membrane-mediated dimerization of PIP5K on supported lipid bilayers. We find that the lifetime of PIP5K dimers is transient relative to the total dwell time of a single membrane bound kinase. Sequence conservation shared between the dimer interfaces of PIP5K paralogs facilitates heterodimerize, which we detected based on changes in smFRET efficiency, single molecule dwell times, and membrane diffusivity. Based on our results, it’s important to consider the total cellular concentration of PIP5K paralog when trying to predict the extent of membrane-mediated dimerization at the cellular plasma membrane. Environmental conditions that modulate gene expression of PIP5K paralogs is predicted to change the total cellular concentration of PI(4,5)P_2_ lipids by enhancing membrane-mediated dimerization. It remains unclear what homeostatic mechanisms cells have evolved to respond to tissue specific changes in PIP5K expression in healthy and disease states.

Several studies have shown that PI(4,5)P_2_ lipid synthesis is inhibited by PIP4K expression (24, 25, 39). This is mediated by a direct interaction between PIP4K and PIP5K, which attenuates PIP5K lipid kinase activity (24, 25). Here, we demonstrate that PIP4K blocks membrane-mediated dimerization of PIP5K, via an interface that is also required to attenuate PIP5K lipid kinase activity in vitro and in vivo. Although we previously reported interaction between PIP4K and PIP5K on supported lipid bilayers using smTIRF microscopy (25), the molecular basis of this interaction was unknown prior to this work. Recent advances in the structural prediction of protein-protein interactions (36) allowed us to model different PIP4K-PIP5K binding interfaces. While some predicted interactions were not necessary for the interaction (i.e. PIP4K R104, **Figure S6**), our mutational analysis confirmed that PIP4K residues M60, L61, and M62 mediate interactions with the dimer interface of PIP5K. Importantly, these mutations did not alter the structural integrity, catalytic activity or intrinsic membrane binding properties of PIP4K. Comparing these mutation to those previously reported to disrupt the interaction with PIP5KA (24), we have identified the same hydrophobic loop as driving the interaction, although the reported mutations in Wang et al are expected to have a less potent effect, with mostly like-for-like hydrophobic substitutions in the core MLM motif and only a single hydrophobic residue, M60, mutated to a charged residue (M60E in that case). Importantly, this hydrophobic motif is conserved even in distantly related eukaryotic clades, such as recently identified choanoflagellate and sponge PIP4K orthologues (40).

PIP5K activity is greatly enhanced during Wnt signaling pathway and phagocytosis (3, 41, 42). How PIP5K activity is enhanced during these processes is not well understood, though enhance dimerization provides a plausible explanation. Dimerization could facilitate interactions with GTPases like Rac1 or Arf6, or by other proteins that have been shown to interact with PIP5K like Daam2 or Dishevelled (29, 43–45). These proteins could also cluster PIP5K into distinct signalosomes on the inner leaflet that promoted density-dependent dimerization. Future studies that attempt to describe PIP5K dimerization in vivo should explore dimerization in terms of cell signaling pathways like Wnt to attempt to describe a functional role for dimerization.

Recent studies have implicated PIP5K activity being necessary for tumor progression in both prostate and breast cancers (46, 47). PI(4,5)P_2_ generated by PIP5K is a precursor for second messengers, including inositol trisphosphate (IP_3_) and diacylglycerol (DAG) generated by phospholipase C (PLC) at the plasma membrane. In addition, PI(4,5)P_2_ lipids are the substrate for PI3K-mediated production of phosphatidylinositol-3,4,5-trisphosphate (PI(3,4,5)P_3_). Elevated levels of PI(3,4,5)P_3_ drive Akt/mTOR pathway activation, promoting undesired cell proliferation and cancer cells metastasis (48–50). Similarly, misregulation of PIP4K increases the total plasma membrane density of PI(4,5)P_2_, which promotes elevated levels of PI(3,4,5)P_3_ following stimulation with insulin (24, 51). Having established that membrane-mediated dimerization of the PIP5K as a node for tuning PI(4,5)P_2_ synthesis, deciphering how various cellular conditions modulate PIP5K dimerization and PIP4K-PIP5K complex formation will be critical for deciphering PIP lipid homeostasis in healthy and diseases states.

## ACKNOWLEDGMENTS

We thank Evan LeGrand and Mike Harms (University of Oregon) for sharing their Jasco J-815 circular dichroism spectrometer and protocols.

## SUPPORTING INFORMATION

This article contains supporting information.

Supplemental Figures S1-S9

Supplemental Movie S1

Plasmid inventory

## AUTHOR CONTRIBUTIONS

Resources: B.R.D., M.J.C., G.R.H., S.D.H.

Experiments and investigation: B.R.D., M.J.C., S.M.F., G.R.H., S.D.H.

Data Analysis: B.R.D., M.J.C., S.M.F., G.R.H., S.D.H.

Conceptualization: B.R.D., G.R.H., S.D.H.

Interpretation: B.R.D., G.R.H., S.D.H.

Data curation: B.R.D., M.J.C., G.R.H., S.D.H.

Writing – Review and editing: B.R.D., M.J.C., S.M.F., G.R.H., S.D.H.

Writing – Original draft: B.R.D., S.D.H.

Supervision: G.R.H., S.D.H.

Project administration: G.R.H., S.D.H.

Funding acquisition: G.R.H., S.D.H.

## DATA AVAILABILITY

All the information needed for interpretation of the data is presented in the manuscript or the supplemental material. Plasmids related to this work are available upon request.

## MATERIALS & METHODS

### Molecular Biology

For recombinant protein expression and purification, the following genes were used for cloning: PH domain derived from human phospholipase C-*δ*1 (PLCδ Accession #P51178.2), human phosphatidylinositol 4-phosphate 5-kinase type-1 beta (hPIP5K1B; Uniprot #O14986) and human phosphatidylinositol 5-phosphate 4-kinase type-2 alpha (hPIP4K2A; Uniprot #P48426). The gene encoding yeast Mss4 (Uniprot #P38994) was obtained by PCR amplification from yeast genomic DNA (S288C, *Saccharomyces cerevisiae*). The gene encoding mouse PIP5KA isoform 1 was purchased as a cDNA clone from Horizon Discovery (cat# MMM1013-202762630, Uniprot #P70182). For simplicity, hPIP4K2A is referred to as PIP4KA throughout the manuscript. PIP5K1A is referred to as PIP5KA. Gene sequences were subcloned into either bacterial, insect cell, or mammalian expression vectors using Gibson assembly. Plasmids containing PIP5KB mutations (i.e. C110S, D51R, D266K, P163C, C411S), PIP4KA (R104D, M60R, L61E, M62R), and PIP5KA (D92R) were generated through site-directed mutagenesis using the PfuUltra High-Fidelity DNA polymerase (Agilent, cat# 600380). The PIP5KB FRET construct contains the following point mutations: C110S, P163C, and C411S. The complete open reading frame of all vectors used in this study were sequenced by Azenta (formerly Genewiz) and Plasmidsaurus (University of Oregon) to ensure constructs lacked deleterious mutations. Each protein expression construct was screened for optimal yield and solubility in either bacteria (BL21 DE3 Star, Rosetta, etc.) or *Spodoptera frugiperda* (Sf9) insect cells. Note that there have previously been inconsistencies in nomenclature between human and mouse PIP5K paralogs. In this manuscript, PIP5KB refers to the human PIP5KB paralog. All plasmids created and used within this manuscript function under this human nomenclature.

Plasmids employed pEGFP-C1 or -N1 backbones (Clontech) or derivatives thereof. Fluorophores included a human codon-optimized EGFP from Aequorea victoria (F64L/S65T; (52), mCherry (a monomeric Discosoma variant; (53), iRFP from Rhodopseudomonas palustris BphP2 (10.1038/nbt.1918), and mTagBFP2 from Entacmaea quadricolor eqFP578 (54). Site-directed mutagenesis was performed with complementary oligos (ThermoFisher). The respective high and low affinity PI(4,5)P_2_ biosensors Tubby_c_^R332H^ and Tubby were described previously (37). Additionally, the EGFP-PIP5K1A, TagBFP2-PIP4KA, Myr-mCherry-PIP4KA and mCherry-FKBP-PIP4KA constructs are as previously described (25), as is the PM-targeted FRB fused to the n-terminal 11 residues of Lyn kinase and iRFP (52). PIP4KA M60R,L61EM62R was generated by PCR with primer pair ccctgttaGAGAgaGAccagatgacttcaaagcctattc and agtcatctggTCtcTCTCtaacagggatttgaacatggctc. Δ54-62→GGSGG mutants were generated by PCR with primer pair GGTGGCTCTGGAGGGgcctattcaaaaataaaggtggac and gctcagttcattgatcgagtgg, which produced the deletion and insertion of the GGSGG linker; the product was phosphorylated, and blunt ends were re-ligated. The A671E mutation was similarly introduced by PCR and blut-end ligation using primer pair GAGcatgctgcaaaaactgttaaacatggcgctg and agctttcttttttgcatcataatgagtaaggatgtc. All constructs were sequence verified by Sanger or next generation sequencing and are available via Addgene (www.addgene.org).

### Rational design of PIP5KB FRET construct

Accessible surface area calculations were performed on zebrafish PIP5KA (4TZU.pdb) and human PIP5KB AlphaFold structure using UCSF Chimera. Following labeling of native solvent exposed cysteines in wild type PIP5KB with AF488-maleimide, the kinase was trypsin digested and analyzed by electrospray ionization mass spectroscopy (ESI-MS) (UC Berkeley MS Facility). Through trial and error, we determined that mutation C175S disrupts kinase activity of recombinantly purified PIP5KB (data not shown). The ideal combination of the mutations for the PIP5KB FRET construct presented here is C110S, P163C, C411S.

### Protein purification

#### PIP5KB

Gene sequences encoding PIP5KB 1-421aa (wild type, D51R, and FRET construct) were cloned into FastBac1 vectors in frame with a N-terminal his_6_-MBP-TEV-GGGGG and expressed under the polyhedrin (pH) promoter. BACMIDS and baculoviruses were generated as previously described (55). For large protein expression, high five cells were infected with baculovirus using an optimized multiplicity of infection (MOI), typically 2% vol/vol. Infected insect cells were grown for 48 hours at 27°C in ESF 921 Serum-Free Insect Cell Culture medium (Expression Systems, Cat# 96-001-01). Cells were then harvested by centrifugation, washed with 1x PBS [pH 7.2], resuspended in cell storage buffer (1x PBS [pH 7.2], 10% glycerol, 2x Sigma protease inhibitor table), and then stored in the −80°C freezer. For purification, frozen insect cell pellets for 2-4 liters of liquid culture were thawed at room temperature in a water bath and lysed into buffer containing 50 mM Na_2_HPO_4_ [pH 8.0], 10 mM imidazole, 400 mM NaCl, 1 mM PMSF (added twice, once before homogenization and once after), 5 mM BME, 100 µg/mL DNase, SIGMAFAST protease inhibitor cocktail tablets, EDTA-free (Sigma, Cat# S8830-20TAB) per 100 mL lysis buffer. Cells in this buffer were lysed using a dounce homogenizer. Lysate was clarified by centrifugation at 36,000 rpm (140,000 x *g*) for 60 minutes under vacuum using a Beckman Ti-45 rotor at 4°C. Lysate was then batch bound to 5 mL of Ni-NTA Agarose (Qiagen, Cat# 30230) resin at 4°C for 2 hours in a beaker set on a stir plate. Resin was then collected in 50 mL tubes, centrifuged, and washed with buffer containing 50 mM Na_2_HPO_4_ [pH 8.0], 10 mM imidazole, 400 mM NaCl, and 5 mM BME and centrifuged again before being transferred to gravity flow column in more wash buffer. Ni-NTA resin with his_6_-MBP-TEV-GGGGG-PIP5K bound was then eluted into buffer containing 500 mM imidazole. Peak fractions were pooled, combined with 200 µg/mL his6-TEV(S291V) protease, and dialyzed against 4 liters of buffer containing 20 mM Tris [pH 8.0], 200 mM NaCl, 2.5 mM BME for 16-18 hours at 4°C. Dialysate was then combined 1:1 with 20 mM Tris [pH 8.0], 1 mM TCEP (~100 mM NaCl final). Precipitation was removed by centrifugation and 0.22 µm syringe filtration. Clarified dialysate was then bound to a MonoS cation exchange column (GE Healthcare, Cat# 17-5168-01) equilibrated in 20 mM Tris [pH 8.0], 100 mM NaCl, 1 mM TCEP buffer. Proteins were resolved over a 10-100% linear gradient (0.1-1 M NaCl, 45 CV, 45 mL total, 1 mL/min flow rate). PIP5K paralogs typically eluted from the MonoS column in the presence of 370-450 mM NaCl. Peak fractions containing PIP5K were pooled, concentrated in a 30 kDa MWCO Vivaspin 6 centrifuge tube (GE Healthcare, Cat# 28-9323-17), and loaded on a 24 mL Superdex 200 10/300 column GE Healthcare, Cat# 17-5174-01) equilibrated in 20 mM Tris [pH 8.0], 200 mM NaCl, 10% glycerol, 1 mM TCEP. Peak fractions were concentrated using a 30 kDa MWCO Vivaspin 6 centrifuge tube and snap frozen at a final concentration of 10-40 µM using liquid nitrogen.

PIP5KB FRET constructs (C110S, P163C, C411S) were fluorescently labeled via their singular surface exposed cysteine residue with either Alexa647-C2-maleimide or Alexa555-C2-maleimide (ThermoScientific). After quenching the labeling reaction with 10 mM DTT, unreacted dye was buffer exchange using a Cytiva NAP-25 column (Fisher scientific, Cat# 10004064) and then loaded on a 24 mL Superdex 200 10/300 column GE Healthcare, Cat# 17-5174-01) equilibrated in 20 mM Tris [pH 8.0], 200 mM NaCl, 10% glycerol, 1 mM TCEP. Peak fractions were concentrated using a 30 kDa MWCO Vivaspin 6 centrifuge tube and snap frozen at a final concentration of 1-30 µM using liquid nitrogen.

#### PIP5KA

Gene sequences encoding PIP5KA (wild type and D92R) were cloned into FastBac1 vectors in frame with an N-terminal his6-MBP-(Asn)_10_-TEV-GGGGG. Protein expression in High Five insect cells was constitutive under the polyhedrin (pH) promoter. High Five cells were infected with baculovirus using an optimized MOI, typically 1.5–2% vol/vol, was determined empirically from small-scale test expression (25–50 mL culture). Infected cells were grown at 27°C for 48 hours in ESF 921 Serum-Free Insect Cell Culture medium (Expression Systems, Cat# 96-001-01). Cells were harvested by centrifugation, washed with 1× PBS (pH 7.2), and then stored in the –80°C freezer. For purification, frozen insect cell pellets for 4–6 L of liquid culture were thawed in an ambient water bath and lysed into buffer containing 50 mM Na_2_HPO_4_ (pH 8.0), 10 mM imidazole, 400 mM NaCl, 1 mM PMSF, 5 mM BME, 100 µg/mL DNase, 1 Sigma protease inhibitor cocktail EDTA free per 100 mL lysis buffer using a dounce homogenizer. Lysate was centrifuged using a Beckman Ti-45 rotor at 4°C at 35,000 rpm (140,000 × *g*) for 60 min under vacuum. Lysate was then batch bound to 5 mL of Ni-NTA Agarose (QIAGEN, Cat# 30230) resin at 4°C for 1–2 hr in a beaker set on a stir plate. Resin was then collected in 50 mL tubes, centrifuged, and washed with buffer containing 50 mM Na_2_HPO_4_ (pH 8.0), 10 mM imidazole, 400 mM NaCl, and 5 mM BME before being transferred to gravity flow column. NiNTA resin with his6-MBP-(Asn)_10_-TEV-GGGGG-PIP5K was then washed with 100 mL of 50 mM Na_2_HPO_4_(pH 8.0), 30 mM imidazole, 400 mM NaCl, and 5 mM BME buffer and then eluted into buffer containing 500 mM imidazole. Peak fractions were pooled, combined with 200 µg/mL his6-TEV(S291V) protease, and dialyzed against 4 L of buffer containing 20 mM Tris (pH 8.0), 200 mM NaCl, 2.5 mM BME for 16–18 hr at 4°C. Dialysate was then combined 1:1 with 20 mM Tris (pH 8.0), 1 mM DTT (~100 mM NaCl final). Precipitation was removed by centrifugation and 0.22 µm syringe filtration. Clarified dialysate was then bound to a MonoS cation-exchange column (GE Healthcare, Cat# 17-5168-01) equilibrated in 20 mM Tris (pH 8.0), 100 mM NaCl, 1 mM TCEP buffer. Proteins were resolved over a 10–100% linear gradient (0.1–1 M NaCl, 45 CV, 45 mL total, 1 mL/min flow rate). PIP5K homologs and paralogs typically eluted from the MonoS column in the presence of 370–450 mM NaCl. Peak fractions containing PIP5K (or Mss4) were pooled, concentrated in a 30 kDa MWCO Vivaspin 6 centrifuge tube (GE Healthcare, Cat# 28-9323-17), and loaded onto a 24 mL Superdex 200 10/300 GL (GE Healthcare, Cat# 17-5174-01) size-exclusion column equilibrated in 20 mM Tris (pH 8.0), 200 mM NaCl, 10% glycerol, 1 mM TCEP. Peak fractions were concentrated in a 30 kDa MWCO Vivaspin 6 centrifuge tube and snap-frozen at a final concentration of 20–40 µM using liquid nitrogen.

#### PIP4KA

Codon optimized gene sequence encoding human PIP4KA (Uniprot # P48426) was cloned into a pETM-derived bacterial expression vector to create the following fusion protein: his_6_-SUMO3-GGGGG-PIP4KA (1-416aa; wild type and mutants). Expressed in BL21(DE3) Star *E. coli* as previously described (25). Bacterial cultures (grown in 2-4 liters of Terrific Broth) were grown at 37°C until OD_600_=0.6. Protein expression was induced with 50 µM IPTG and bacteria expressed protein for 20 hours at 18°C before being harvested by centrifugation. For purification, cells were lysed into buffer containing 50 mM Na_2_HPO_4_ [pH 8.0], 400 mM NaCl, 0.4 mM BME, 1 mM PMSF, DNase, 1 mg/mL lysozyme using a microtip sonicator. Lysate was centrifuged at 16,000 rpm (35,172 x *g*) for 60 minutes in a Beckman JA-17 rotor chilled to 4°C. Lysate was circulated over 5 mL HiTrap Chelating column (GE Healthcare, Cat# 17-0409-01) that had been equilibrated with 100 mM CoCl_2_ for 1 hour, washed with MilliQ water, and followed by buffer containing 50 mM Na_2_HPO_4_ [pH 8.0], 400 mM NaCl, 0.4 mM BME. Recombinant PIP4KA was eluted with a linear gradient of imidazole (0-500 mM, 8 CV, 40 mL total, 2 mL/ min flow rate). Peak fractions were pooled, combined with 50 µg/mL of his6-SenP2 (SUMO protease), and dialyzed against 4 liters of buffer containing 25 mM Na_2_HPO_4_ [pH 8.0], 400 mM NaCl, and 0.4 mM BME for 16-18 hours at 4°C. Following overnight cleavage of the SUMO3 tag, dialysate containing his6-SUMO3, his6-SenP2, and GGGGG-PIP4KA was recirculated for at least 1-2 hr over a 5 mL HiTrap(Co^+2^) chelating column. Flow-through containing GGGGG-PIP4KA was then concentrated in a 30 kDa MWCO Vivaspin 6 before loading onto a Superdex 200 size exclusion column equilibrated in 20 mM HEPES [pH 7], 200 mM NaCl, 10% glycerol, 1 mM TCEP. Peak fractions collected from the Superdex 200 were concentrated in a 30 kDa MWCO Vivaspin 6 centrifuge tube and snap frozen at a final concentration of 20-80 µM using liquid nitrogen.

#### PLCδ-PH and LactC2

Human PLCδ-PH domain (11-140aa) was expressed in BL21 (DE3) Star *E. coli* as a his_6_-SUMO3-(Gly)_5_-PLCδ (11-140aa) fusion protein as previously described (55). The phosphatidylserine lipid sensor, LactC2, was expressed as a his6-TEV-SUMO3-GGGG-LactC2 (271-427aa) fusion protein as previously described (56). When cloning LactC2 for recombinant protein expression, we incorporated a single cysteine at the C-terminal position of the LactC2 open reading frame. Following growth at 37°C in Terrific Broth to an OD_600_ of 0.8, bacterial cultures were shifted to 18°C for 1 hour, induced with 0.1 mM IPTG, and allowed to express protein for 20 hours at 18°C before being harvested. Cells were lysed into 50 mM Na_2_HPO_4_ [pH 8.0], 300 mM NaCl, 0.4 mM BME, 1 mM PMSF, 100 µg/mL Dnase using a microfluidizer. Lysate was then centrifuged at 16,000 rpm (35,172 x *g*) for 60 minutes in a Beckman JA-17 rotor chilled to 4°C. Lysate was circulated over 5 mL HiTrap Chelating column (GE Healthcare, Cat# 17-0409-01) charged with 100 mM CoCl_2_ for 1 hour. Bound protein was then eluted with a linear gradient of imidazole (0-500 mM, 8 CV, 40 mL total, 2 mL/min flow rate). Peak fractions were pooled, combined with SUMO protease (50 µg/mL final concentration), and dialyzed against 4 liters of buffer containing 50 mM Na_2_HPO_4_ [pH 8.0], 300 mM NaCl, and 0.4 mM BME for 16-18 hours at 4°C. Dialysate containing SUMO cleaved protein was recirculated for 1 hr over a 5 mL HiTrap Chelating column. Flow-through containing (Gly)_5_-PLCδ (11-140aa) was then concentrated in a 5 kDa MWCO Vivaspin 20 before being loaded on a Superdex 75 size exclusion column equilibrated in 20 mM Tris [pH 8.0], 200 mM NaCl, 10% glycerol, 1 mM TCEP. Peak fractions containing (Gly)_5_-PLCδ (11-140aa) were pooled and concentrated to a concentration of 75 µM before snap freezing with liquid nitrogen and storage at −80°C. Peak fractions containing (Gly)_5_-LactC2 (271-427aa) were pooled and concentrated to a concentration of 123 µM before snap freezing with liquid nitrogen and storage at −80°C. Biophysical parameters of GGGG-PLC⊠: pI = 7.19, ε_280_ = 17,990 M^-1^•cm^-1^, 15.8 kDa. Biophysical parameters of GGGG-LactC2: pI = 8.99, ε_280_ = 44,920 M^-1^•cm^-1^, 18.1 kDa. As previously described (55), GGGGG-PLCδ (11-140aa) was labeled with either AF488 or Cy3 using sortase mediated peptide ligation. LactC2 was labeled with maleimide DY-647 (Dyomics) using methods previously described (57).

#### Circular dichroism

(CD) experiments were performed using a Jasco J-815 CD spectrometer. Samples containing 15 µM of either PIP4KA or PIP4K (M60R, L61E, M62R) in 20 mM HEPES [pH 7.5], 150 mM NaCl, 5% glycerol, 1 mM TCEP buffer were loaded into a quartz cuvette with a 1 mm pathlength. CD spectra were collected using the Jasco Spectra Manager software at scan speed of 10 nm per minute, standard sensitivity, and 1 sec digital integration time. Spectral scans were collected across the wavelengths of 180-300 nm in triplicates at room temperature (23°C). Background subtraction of CD spectra shown in Figure S7 was accomplished by scanning the buffer alone, which contained 20 mM HEPES [pH 7.5], 150 mM NaCl, 5% glycerol, 1 mM TCEP. Background scans (n = 3) were averaged, and the signal was subtracted from the data plotted in Figure S7. Note that signal measured below 200 nm is noisy because the interaction between circularly polarized light and Cl^−^ ions in our buffer.

#### Size exclusion chromatography (SEC)

To compare the relative molecular weights and conformations of PIP4KA or PIP4K (M60R, L61E, M62R) in solution, a 0.5 mL sample of 7.5 µM purified protein was loaded on 24 mL Superdex200 GL 10/300 SEC column (GE Healthcare, cat# 17-5174-01). The column was equilibrated in buffer containing 20 mM HEPES [pH 7.5], 150 mM NaCl, 1 mM TCEP and run at 0.6 mL/min at 4°C. In parallel, a BioRad molecular weight standard (BioRad, cat# 1511901) containing 5 mg total protein was injected and resolved. The MW standard contained species of the following molecular weights: 1.35, 17, 44, 158, 670 kDa.

#### Preparation of small unilamellar vesicles

The following lipids were used to generated small unilamellar vesicles (SUVs): 1,2-dioleoyl-sn-glycero-3-phosphocholine (18:1 DOPC, Avanti # 850375C), 1,2-dioleoyl-sn-glycero-3-phospho(1’-myo-inositol-4’-phosphate) (18:1 PI(4)P, Avanti Cat# 850151P), L-α-phosphatidylinositol-4-phosphate (Brain PI(4)P, Avanti Cat# 840045X), L-α-phosphatidylinositol-4,5-bisphosphate (Brain PI(4,5)P_2_, Avanti Cat# 840046X). To make liposomes, 2 µmoles total lipids are combined in a 35 mL glass round bottom flask with 2 mL of chloroform. Lipids were dried to a thin film using rotary evaporation with the glass round-bottom flask submerged in a 42°C water bath. The lipid film was then resuspended in 2 mL of PBS [pH 7.2], getting a final concentration of 1 mM total lipids. All lipid mixtures expressed as percentages (e.g. 98% DOPC, 2% PI(4)P) are equivalent to molar fractions. To generate SUVs, 1 mM total lipid mixture was extruded through a 0.05 µm pore size 19 mm polycarbonate membrane (Avanti, Cat# 610002) with filter supports (Avanti, Cat# 610014) on both sides of the polycarbonate membrane. Lipids were then extruded 11 times total to achieve ~50nm SUV size.

#### Preparation of supported lipid bilayers

25×75 mm coverglass (IBIDI, Cat# 10812) was used to create supported lipid bilayers. To clean the coverglass before use, we submerged glass slides in 2% Hellmanex III (Fisher, Cat# 14-385-864) heated to 60-70°C. Coverslides were incubated in heated Hellmanex in a glass coplin jar for at least 30 minutes. The coverglass was then rigorously washed with MilliQ water to finish pre-cleaning. Coverglass was then etched in Pirahna solution (1:3, hydrogen peroxide:sulfuric acid) for 10-15 minutes the same day SLBs were formed. The etched coverglass was again thoroughly rinsed with MilliQ water before being rapidly dried with nitrogen glass. Once dried, glass was adhered to a 6-well sticky-side chamber (IBIDI, Cat# 80608). SLBs were formed by flowing 30 nm SUVs diluted in PBS [pH 7.2] to a total lipid concentration of 0.25 mM and incubated for 30 minutes. IBIDI chambers were then washed with 5 mL of PBS [pH 7.2] to remove non-absorbed SUVs. Membrane defects are blocked for 15 minutes with a 1 mg/mL beta casein (ThermoFisher, Cat# 37528) diluted in 1x PBS [pH 7.4]. Before use as a blocking protein, frozen 10 mg/mL beta casein stocks were thawed, centrifuged for 30 minutes at 21370 x *g*, and 0.22 µm syringe filtered. After blocking SLBs with beta casein, membranes were washed again with 1mL of PBS, followed by 1 mL of kinase buffer before TIRF-M.

#### Assay for measuring the kinetics of PI(4,5)P_2_ production

The kinetics of PI(4)P phosphorylation were measured on SLBs formed in IBIDI chambers and visualized using TIRF microscopy as previously described. Imaging or reaction buffer contained 20 mM HEPES [pH 7.0], 150 mM NaCl, 1 mM ATP, 5 mM MgCl_2_, 0.5 mM EGTA, 20 mM glucose, 200 µg/mL beta casein (ThermoScientific, Cat# 37528), 20 mM BME, 320 µg/mL glucose oxidase (Serva, Cat# 22780.01 *Aspergillus niger*), 50 µg/mL catalase (Sigma, Cat# C40-100MG Bovine Liver), and 2 mM Trolox (UV treated as previously described by Hansen et al. 2019). Perishable reagents (i.e. glucose oxidase, catalase, and Trolox) were added 5-10 minutes before image acquisition. For all experiments, we monitored the change in PI(4,5)P_2_ membrane density using a solution concentration of 20 nM AF488-GGGGG-PLCδ, Cy3-GGGGG-PLCδ, or AF647-GGGGG-PLCδ. Density of PIP lipids (lipids/µm^2^) was calculated assuming a footprint of 0.72 nm^2^ for DOPC lipids (55, 58).

#### Quantification of lipid kinase activity

When comparing PIP5K lipid kinase activity across conditions (+/– PIP4K), we report the time to 95% reaction completion. Since PIP4K-mediated inhibition of PIP5K does not occur until a threshold density of PI(4,5)P_2_ is generated to efficiently recruit PIP4K, there is a delay in the membrane recruitment of PIP4K. Therefore, comparing the reaction half-time would yield similar PIP5K reaction kinetics in absence or presence of PIP4K.

#### Microscope hardware and imaging acquisition

Single molecule imaging experiments were performed on an inverted Nikon Ti2 microscope using a 100x Nikon objective (1.49 NA) oil immersion TIRF objective. The x-axis and y-axis positions were manually controlled using a Nikon motorized stage and joystick. All images were acquired using an iXion Life 897 EMCCD camera (Andor Technology Ltd., UK). Fluorescently labeled proteins were excited with either a 405 nm, 488 nm, 561 nm, or 637 nm diode laser (OBIS laser diode, Coherent Inc. Santa Clara, CA) controlled by a Vortran laser drive with acousto-optic tunable filters (AOTF) control. The power output measured through the objective for single particle imaging was 1-3 mW. For dual color imaging of mNG-PIP5K localization during Cy3-PLCδ monitored PI(4,5)P_2_ synthesis, samples were excited with 1 mW 488 nm and 1 mW 561 nm light, as measured through the objective. Excitation light was passed through the following dichroic filter cubes before illuminating the sample: (1) ZT488/647rpc and (2) ZT561rdc (ET575LP) (Semrock). Fluorescence emission was detected on an ANDOR EMCCD camera position after a Sutter emission filter wheel housing the following emission filters: ET525/50M, ET600/50M, ET700/75M (Semrock). All in vitro experiments were performed at room temperature (23°C). Microscope hardware was controlled using Nikon NIS elements.

#### smFRET imaging acquisition

Single molecule FRET imaging experiments were performed on the same microscope as the previous section described. Membrane-mediated dimerization of PIP5KB was detected based on the sensitize emission of AF647-PIP5KB (acceptor) in the presence of AF555-PIP5KB (donor). Fluorescently labeled proteins were excited with either 532 or 561 nm diode laser (OBIS laser diode, Coherent Inc. Santa Clara, CA) controlled by a Vortran laser drive with acousto-optic tunable filters (AOTF) control. Conditions were optimized to minimal direct excitation of AF647-PIP5KB and channel bleed-through cause by the presence of AF555-PIP5KB. The power output measured through the objective for single particle imaging was 1-3 mW. Excitation light was passed through a ZT561rdc (ET575LP) (Semrock) dichroic filter cubes to illuminate the sample. Fluorescence emission was detected on an ANDOR EMCCD camera position after passing through a Sutter emission filter wheel loaded with the ET700/75M (Semrock) filter. All in vitro experiments were performed at room temperature (23°C). Microscope hardware was controlled using Nikon NIS elements..

#### Image analysis, curve fitting, and statistics for smFRET

Single AF647 labeled PIP5K molecules exhibiting FRET based on sensitized emission were identified using the ImageJ/Fiji Trackmate Plugin (also see *Single Particle Tracking*) (59). By comparing the molecular brightness of particles identified in our controls to our experimental FRET condition, we identified PIP5KB based on a threshold molecular brightness of the acceptor emission that far exceeded the brightest particles in our control conditions. Note that the threshold intensity used to identified particles was set below the threshold intensity used to calculate the frequency of particles that exhibit smFRET. As result some particles that represent low intensity background signal were detected as can be seen in the molecule brightness histograms (e.g. Figure 1G). To plot the frequency of particles observed by smFRET, we calculated the frequency of smFRET particle brightness. This was achieved by dividing the total number of molecules brighter than 99% of particles in the donor and acceptor alone controls by the total number of tracked particles (Frequency = #bright molecules/#total molecules). The number of technical replicates used for each plot is stated in the figure legend (n=3-10). Statistical analysis between the mean of two populations was conducted via GraphPad using an unpaired Student’s T-test with Welch’s correction if variance were unequal.

#### Cell Culture and Transfection

HEK293A cells (ThermoFisher, cat# R70507) were maintained in low-glucose DMEM (ThermoFisher, Cat# 10567022) with 10% heat-inactivated FBS (ThermoFisher, cat# 10438-034), 100 U/mL Penicillin-Streptomycin (ThermoFisher, Cat# 15140122), and 0.1% chemically defined lipid supplement (ThermoFisher, cat# 11905031), at 37 °C and 5% CO_2_. Cells were passaged 1:5 twice weekly using TrypLE (ThermoFisher, cat# 12604039). For imaging, cells were seeded on 35 mm #1.5 glass-bottom dishes (CellVis, cat# D35-20-1.5-N) pre-coated with 5 µg human fibronectin (ThermoFisher, cat# 33016-015). At 50–80% confluency, cells were transfected using 1 µg DNA with 3 µg Lipofectamine2000 (ThermoFisher, cat# 11668019) in 200 µL Opti-MEM (ThermoFisher, cat# 51985091), following the manufacturer’s protocol. Imaging was performed 18–24 hr post-transfection.

#### Live cell confocal microscopy

Prior to imaging, cells were switched to Fluorobrite DMEM (ThermoFisher, cat# A1896702) supplemented with 10% heat-inactivated FBS, 25 mM HEPES (pH 7.4), 2 mM GlutaMAX (ThermoFisher, cat# 35050061), and 0.1% lipid supplement. Media volume was adjusted to 2 mL total after any additions. Imaging was performed using a Nikon TiE A1R confocal system in resonant mode with a 100X 1.45 NA Plan-Apochromat objective. Air temperature was maintained at 37°C by a heated Perspex enclosure around the stage (Tokai Hit). Signal-to-noise was enhanced via 8× frame averaging. Fluorophores were excited using a dual fiber-coupled LUN-V laser launch (405, 488, 561, 640 nm), and emission was collected with Chroma dual pass filters: 420–480/570–620 nm and 505–550/650–850 nm. The pinhole was set to 1.2× the Airy unit of the longest emission wavelength. All images were acquired with Nikon Elements and saved as .nd2 files.

#### Image Analysis

Images were analyzed in Fiji (60). A custom macro was used to generate channel-specific montages displaying all x-y positions in a given experiment ROIs were drawn manually on these projections. Fluorescence ratios between compartments were quantified as described (61), using a binary mask generated via à trous wavelet decomposition. Intensity values were normalized to mean ROI pixel intensity to control for variable expression.

#### Single particle tracking

Fluorescent protein detection and tracking was performed using the ImageJ/Fiji TrackMate plugin. Data in the form of .nd2 files were loaded in ImageJ. Before being analyzed using TrackMate, data brightness was adjusted for molecules to be easily identifiable. TrackMate was then used to identify and track molecular tracks in these steps: Particles were first identified using the LoG detector option based on brightness and signal-to-noise ratio. Once identified, particles were tracked for their full lifetime using the LAP tracker. This LAP tracker follows molecular displacement as a function of time. Particle trajectories were filtered based on Track Start (removed trajectories that began in first frame), Track End (removed trajectories present in last frame), Duration (removed trajectories ≤ 2 frames and singular extra-long tracks), Track displacement (removed immobilized particles displacement <0.1), and X - Y location (removed particles near the edge of the images). The TrackMate output files were analyzed using PRISM 9 (GraphPad) to calculate characteristic dwell times and diffusion coefficients.

Step size distributions derived from single particle trajectories were plotted in Prism as frequency versus step size (µm). For all analysis presented in this manuscript, the bin size for the step size distribution equals 0.01 µm. For curving fitting, the step-size distributions were plotted as probability density versus step size (µm). This was achieved by dividing the frequency distribution (i.e. y-axis values) by the bin size (0.01 µm). The probability density versus step size plots were fit to the following one- or two-species distributions:

Single species model:

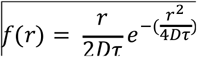

Two species model:

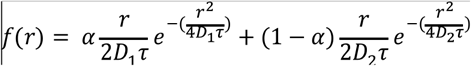

D1=species 1diffusion coefficient (µm^2^/sec), D2= species 2 diffusion coefficient (µm^2^/sec), alpha (α_1_ = % of species 1, r = step size (µm),α_2_ = time interval between steps (sec). Final step size distribution plots were generated in PRISM graphing software and using the following equations: (1 species model): f(x) = x/(2*D1*t)*exp(-(x^2/ (4*D1*t))), (2 species model): f(x) = alpha*(x/(2*D1*t)*exp(-(x^2/(4*D1*t))))+(1-alpha)*(x/(2*D2*t)*exp(-(x^2/ (4*D2*t)))).

To calculate the single molecule dwell times for, FRET-imaged AF647-PIP5KB (C110S, P163C, C411S), standard single molecule imagining of AF647-PIP5KB (and mutants), and AF647-PIP5KA (and mutants) we generated a cumulative distribution frequency (CDF) plot using the frame interval as the bin size (e.g. 50 ms). The log_10_(1-CDF) was plotted against the dwell time and fit to either a single or double exponential decay curve.

Single exponential model:

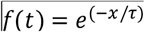

Two exponential model:

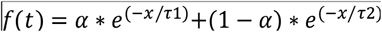

Fitting procedure initiated with a single exponential. In cases of a low-quality single exponential fit, a maximum of two species model was used. For double exponential fit, alpha (α) represents the fraction of fast dissociating population of molecules characterized by the time constant, α_1_. Plots, curve fitting, and statistical analysis of single molecule dwell time and step size distributions were performing using Prism 10 (GraphPad). Single molecule dwell time and step size presented in this manuscript represent combined data from 3 technical replicates with ≥3 movies acquired from multiple fields of view for each experimental condition. Dwell time distributions and curve fits were generated with n = 1000-3000 particle trajectories. Step size distribution plots and curve fits represent 10,000-30,000 measured displacements.

Graphing and statistical analysis were performed in Prism 10 (GraphPad). Representative images were selected for typical morphology, signal-to-noise ratio, and fluorescence levels near the cohort median. Brightness and contrast adjustments were applied uniformly across the entire image if needed.

