## Supplemental Figures for "PIP4K attenuates PIP5K lipid kinase activity by disrupting membrane-mediated dimerization"

#### Supplemental Figure 1

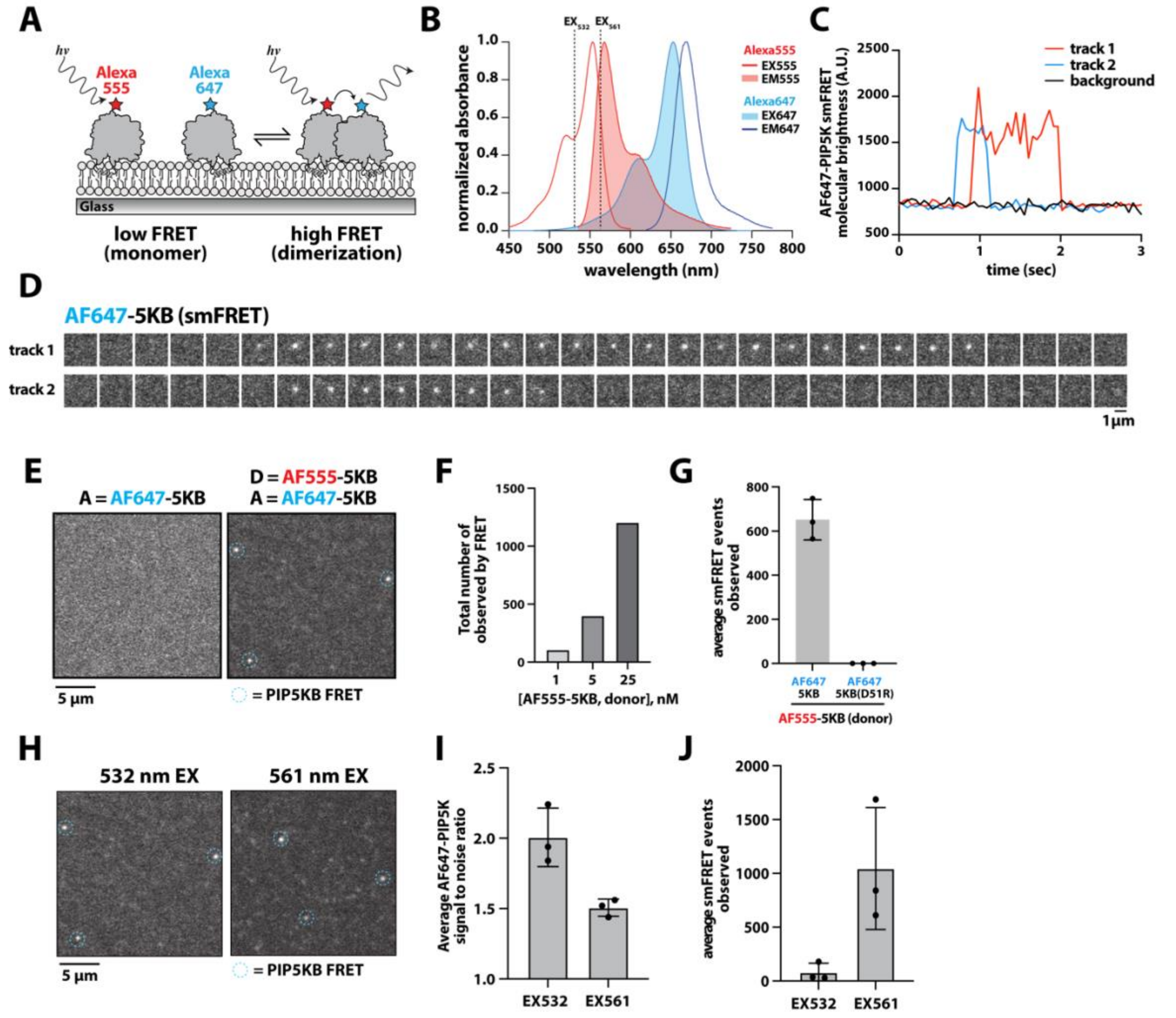


**Supplemental Figure 1**

**Membrane-mediated dimerization of PIP5K visualized by smFRET**

**(A)** Cartoon schematic showing the molecular basis of smFRET based on membrane-mediated dimerization of PIP5KB. **(B)** Excitation-emission spectra for AF555 and AF647. Dashed lines indicate the laser excitation wavelengths (i.e. 532 and 561 nm). Note that 561 can directly excite the acceptor at a low probability. **(C)** Plot showing the fluorescence intensity of AF647-PIP5KB visualized by smFRET TIRF-M. **(D)** Montage of smFRET images for plot in C and Movie S1. Frames separated by 52 ms. **(E)** Representative TIRF-M images showing AF647-PIP5KB visualized by smFRET in the presence of acceptor alone (1 nM AF647-PIP5KB) or donor + acceptor (5 nM AF555-PIP5KB + 1 nM AF647-PIP5KB). **(F)** The total number of AF647-PIP5KB molecules observed increases based on the total concentration of AF555-PIP5KB (donor). **(G)** smFRET requires an intact dimer interface. The acceptor, 1 nM AF647-PIP5K (D51R), does not exhibit FRET in the presence of 5 nM AF555-PIP5KB (donor). **(H)** Representative TIRF-M images showing AF647-PIP5KB visualized by smFRET with either a 532 or 561 nm excitation laser. **(I)** Average signal to noise ratio of AF647-PIP5KB molecules detected by smFRET is greater when excited with a 532 nm laser. **(J)** More AF647-PIP5KB tracks were detected by smFRET when using a 561 nm laser for excitation. **(B-J)** Unless stated otherwise, all experiments include 1 nM AF647-PIP5K plus 5 nM AF555-PIP5KB. The total number of smFRET events were quantified over 30 second period of image acquisition on a membrane surface of 3000 µm$^2$. Membrane composition: 4% PI(4,5)$P_2$, 96% DOPC. Bars equal mean. Errors equal SD from 3 technical replicates.

### Supplemental Figure 2

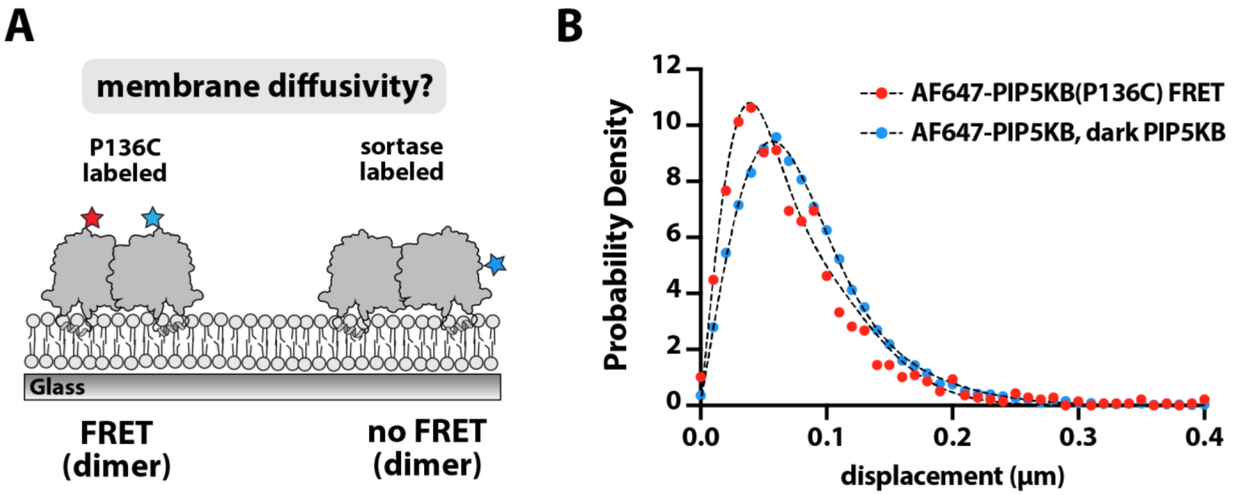


**Supplemental Figure 2**

**Quantification of AF647-PIP5K membrane diffusion visualized by smFRET**

**(A)** Cartoon schematic showing membrane bound PIP5K dimers with different fluorescent labels. Membrane diffusion was measured by monitoring either AF647-PIP5KB(P136C) by smFRET or AF647-PIP5KB labeled via sortase mediated peptide ligation in the presence of 50 nM dark unlabeled PIP5KB. **(B)** Step size distributions showing the displacement of the indicated PIP5K construct. Dashed lines represent the distribution fits with a two species model, which yielded the following diffusion coeffcients: AF647-PIP5KB visualized by smFRET (D1 = 0.01 $\mu m^2$/sec, D2 = 0.048 $\mu m^2$/sec, $\alpha$ = 0.36) and sortase labeled AF647-PIP5KB in the presence of 50 nM dark unlabeled PIP5KB (D1 = 0.027 $\mu m^2$/sec, D2 = 0.084 $\mu m^2$/sec, $\alpha$ = 0.6). Alpha ($\alpha$) equals the fraction of molecules with characteristic diffusion coefficient, D1. Distributions contain ≥10,000 single molecule displacement events. Membrane composition: 4% $PI(4,5)P_2$, 96% DOPC.

### Supplemental Figure 3

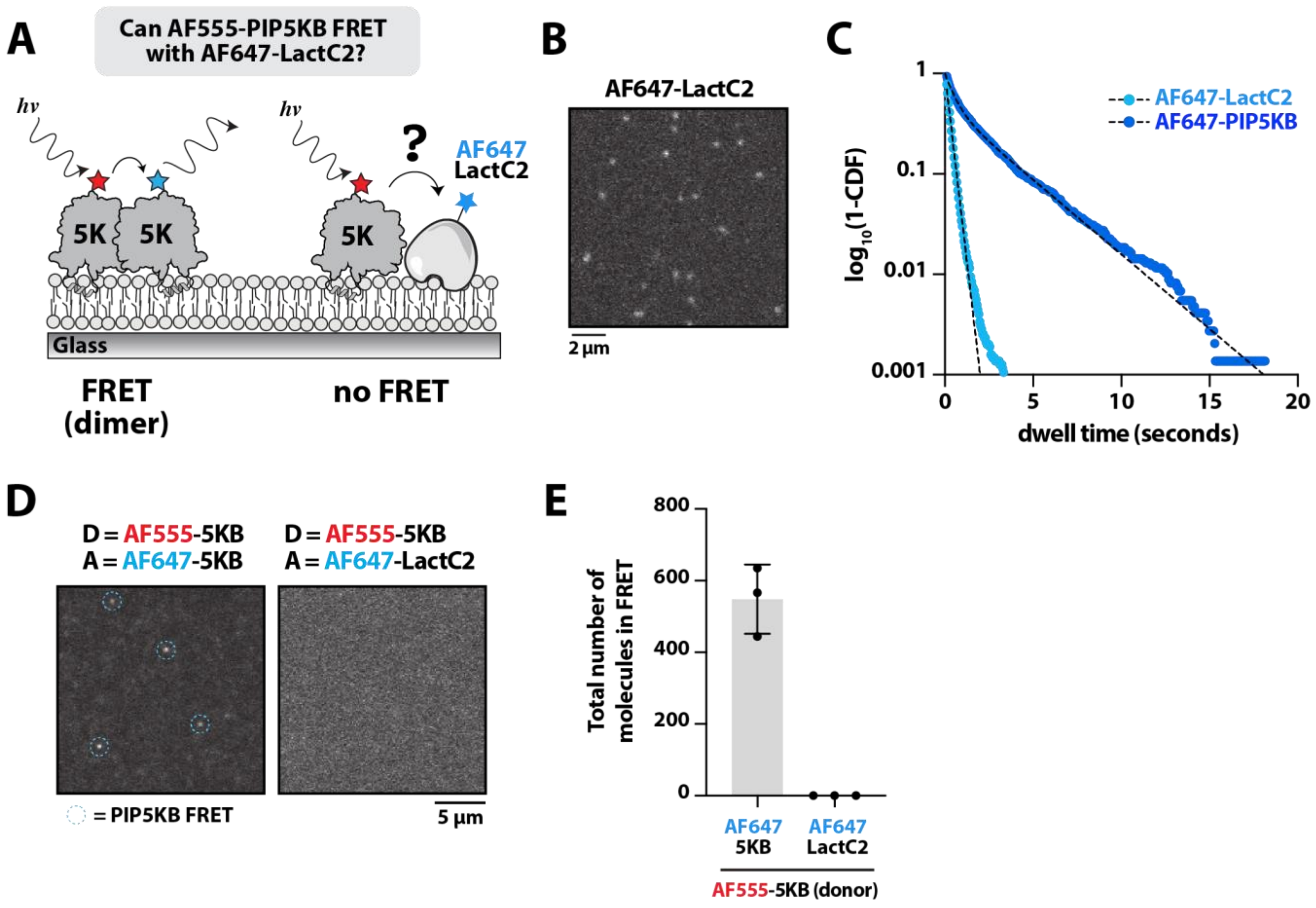


**Supplemental Figure 3**

**PIP5K does not exhibit smFRET in the presence of the peripheral membrane binding protein, LactC2**

**(A)** Cartoon schematic illustrating smFRET based on dimerization between AF555-PIP5KB and AF647-PIP5KB versus non-specific smFRET between AF555-PIP5KB and membrane bound AF647-LactC2. **(B)** Representative single molecule TIRF-M image in the presence of 1 pM AF647-LactC2. Note that LactC2 is best known for its ability to interact with phosphatidylserine (Yeung et al., 2008). In our hands, AF647-LactC2 also displays some specificity for PI(4,5)$P_2$ lipids **(C)** Single molecule dwell time distributions measured in the presence of either 1 pM AF647-LactC2 ($\tau_1$ = 0.289 sec, n = 9488 molecules) or 10 pM AF647-PIP5KB ($\tau_1$ = 0.55 sec, $\tau_2$ = 2.93 sec, n = 1464 molecules). Dwell times were calculated by fitting $\log_{10}$(1-cumulative distribution frequency (CDF)) to a single or double exponential decay curve (black dashed lines). Alpha ($\alpha$) equals the fraction of molecules with the time constant, $\tau_1$. **(D)** AF555-PIP5KB does not exhibit smFRET in the presence of AF647-LactC2. Representative TIRF-M images showing donor emission measured in the presence of 5 nM AF555-PIP5KB plus either 1 nM AF647-PIP5KB or 1 nM AF647-LactC2. **(E)** Quantification of the smFRET frequency for experiment in (D). Bars equal mean. Errors equal SD from 3 technical replicates. **(B-E)** Membrane composition: 4% PI(4,5)$P_2$, 96% DOPC.

### Supplemental Figure 4

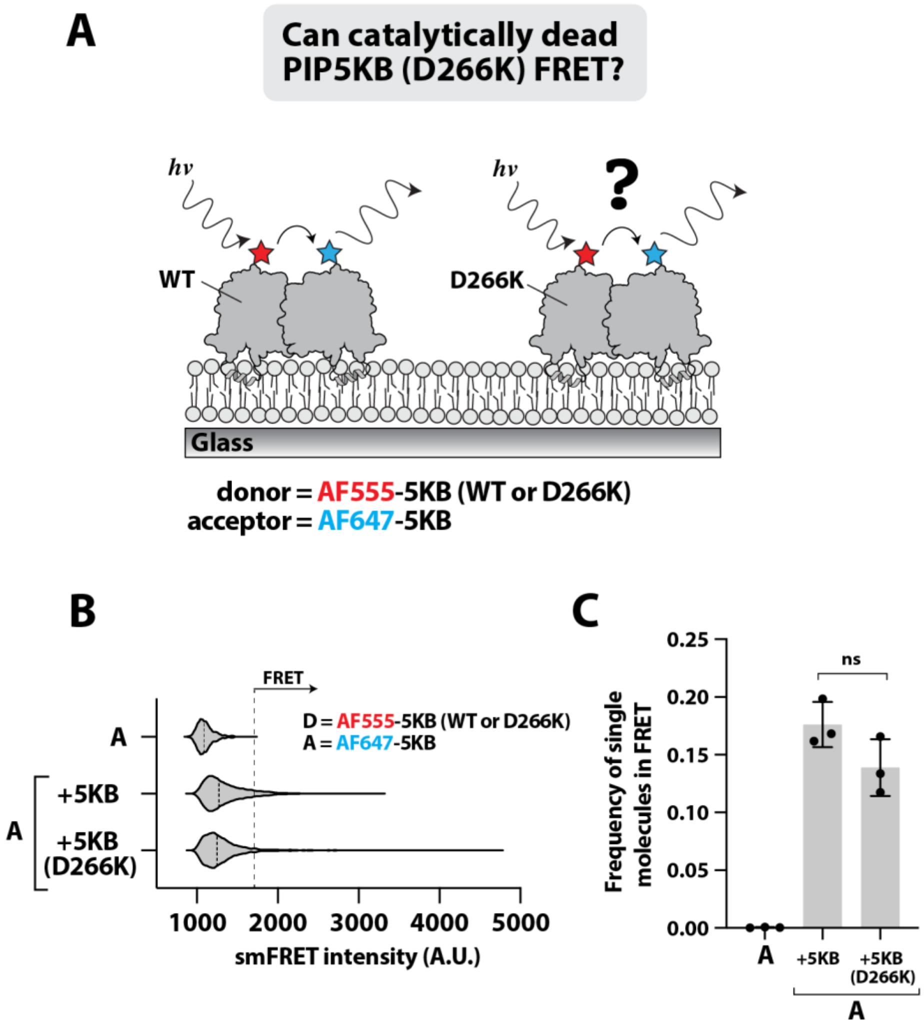


**Supplemental Figure 4**

**Catalytically dead PIP5KB exhibits smFRET based membrane-mediated dimerization**

**(A)** Cartoon schematic showing the molecular basis of smFRET based on membrane-mediated dimerization of PIP5KB, wild type or catalytically dead mutant (D266K). **(B)** smFRET intensity distributions based on the fluorescence emission of AF647-PIP5KB (acceptor). Data collected in the presence 1 nM AF647-PIP5KB FRET acceptor plus 5 nM AF555-PIP5KB FRET donor (wild type or D266K). Dashed line indicates threshold intensity used to quantify frequency of smFRET. **(C)** Frequency of observing PIP5KB dimers by smFRET under the indicated conditions (*p=0.1127* from Student t-test; ns = not significant). Bars equal mean. Errors equal SD from 3 technical replicates. **(B-C)** Membrane composition: 4% $PI(4,5)P_2$, 96% DOPC.

### Supplemental Figure 5

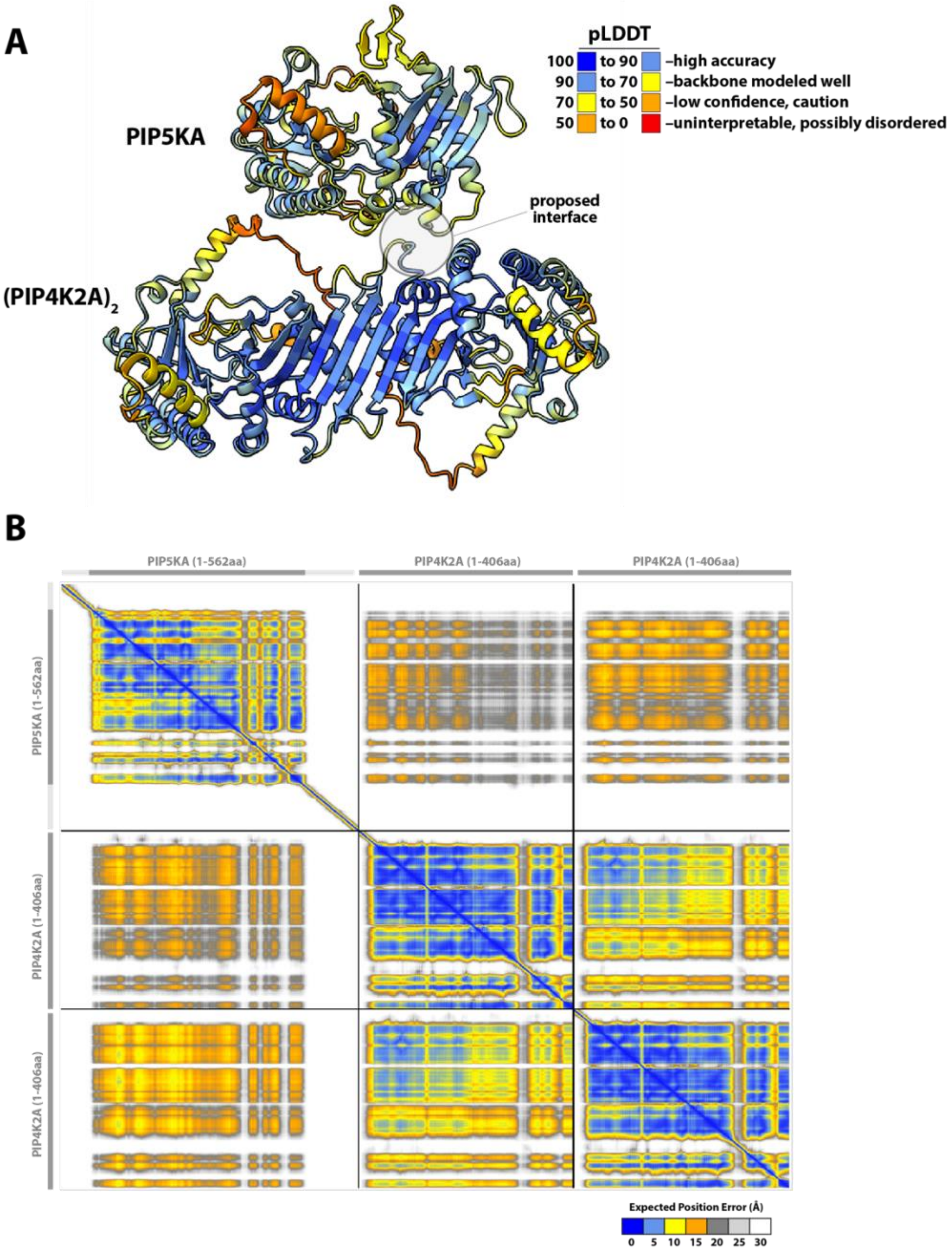


**Supplemental Figure 5**

**Statistical metrics for evaluating the quality of PIP5KA-$(PIP4KA)_2$ AlphaFold prediction**

**(A)** AlphaFold2 multimer prediction of human PIP5KA (1-562 aa) bound to a PIP4K2A (1-406 aa) dimer. For simplicity, only residues corresponding to the kinase domains (i.e. PIP5KA (62-452 aa) and PIP4K2A (36-406 aa)) are shown. The exact amino acid sequences can be found at Uniprot using the following identifiers: hPIP5K1A (Q99755) and hPIP4K2A (P48426). The model is colored by predicted local distance difference (pLDDT) to show regions with high confidence (pLDDT > 90) to low confidence (pLDDT <50). **(B)** Predicted alignment error (PAE) for Alphafold2 multimer model in (A). Note that the PAE plot is not an inter-residue distance map or a contact map. Instead, the coloring indicates expected distance error. The color at (x, y) corresponds to the expected distance error in residue x's position (Angstroms), when the prediction are aligned on residue y (more information can be found at https://alphafold.ebi.ac.uk/).

### Supplemental Figure 6

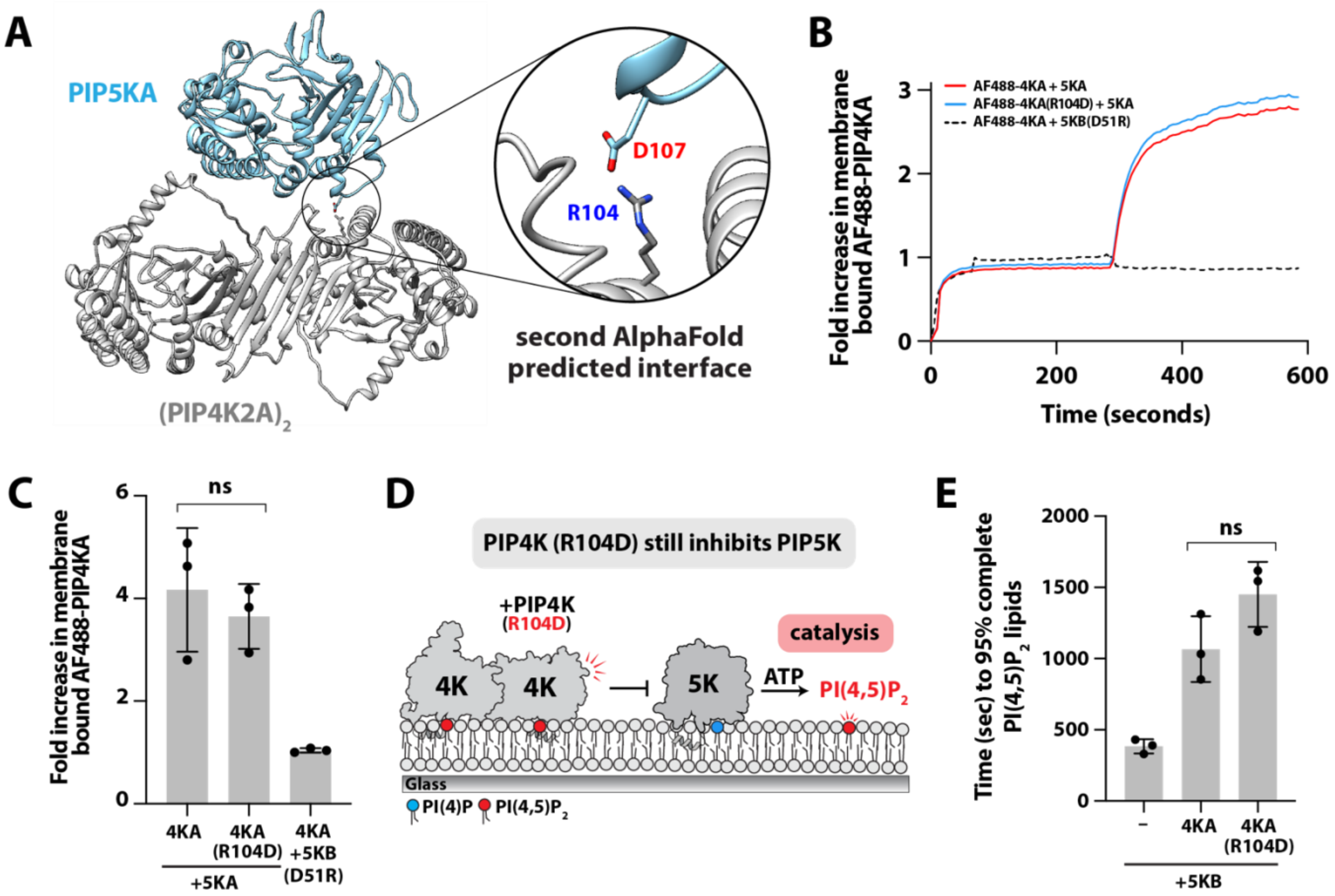


**Supplemental Figure 6**

**The PIP4K (R104R) mutation is not sufficient to disrupt interaction with PIP5KA or PIP5KB**

**(A)** AlphaFold multimer structure prediction of PIP5KA (62-452aa) and $(PIP4KA)_2$ (36-406aa) complex mediated by a salt bridge between residue D107 of human PIP5KA and R104 of human PIP4KA. The homologous residue is D51 in human PIP5KB and D92 in mouse PIP5KA. **(B)** Representative fold-change of 50 nM AF488-PIP4KA (wild type or R104D) membrane binding in the presence of 10 nM unlabeled PIP5KA (dark red) or 10 nM unlabeled PIP5KB (D51R). Membrane composition: 2% $PI(4,5)P_2$, 98% DOPC. **(C)** Fold change in membrane binding quantified from (B). Bars equal mean. Errors equal SD from 3 technical replicates (*p=0.556* Student t-test; ns = not significant). **(D)** Cartoon schematic showing SLB assay for measuring PIP4KA (R104D)-mediated inhibition of PIP5K lipid kinase activity. **(E)** PIP4KA (R104D) does not attenuates PIP5KB activity. Quantification of time required for PIP5KB to generate 95% of maximum $PI(4,5)P_2$ lipid density. Kinase activity measured in the presence of 1 nM PIP5KB +/- 50 nM PIP4KA (wild type or R104D). Reaction monitored in the presence of 20 nM Cy3-PLCδ PH domain. Bars equal mean. Errors equal SD from 3 technical replicates (*p=0.109* Student t-test; ns = not significant). Membrane composition: 4% PI(4)P, 96% DOPC.

### Supplemental Figure 7

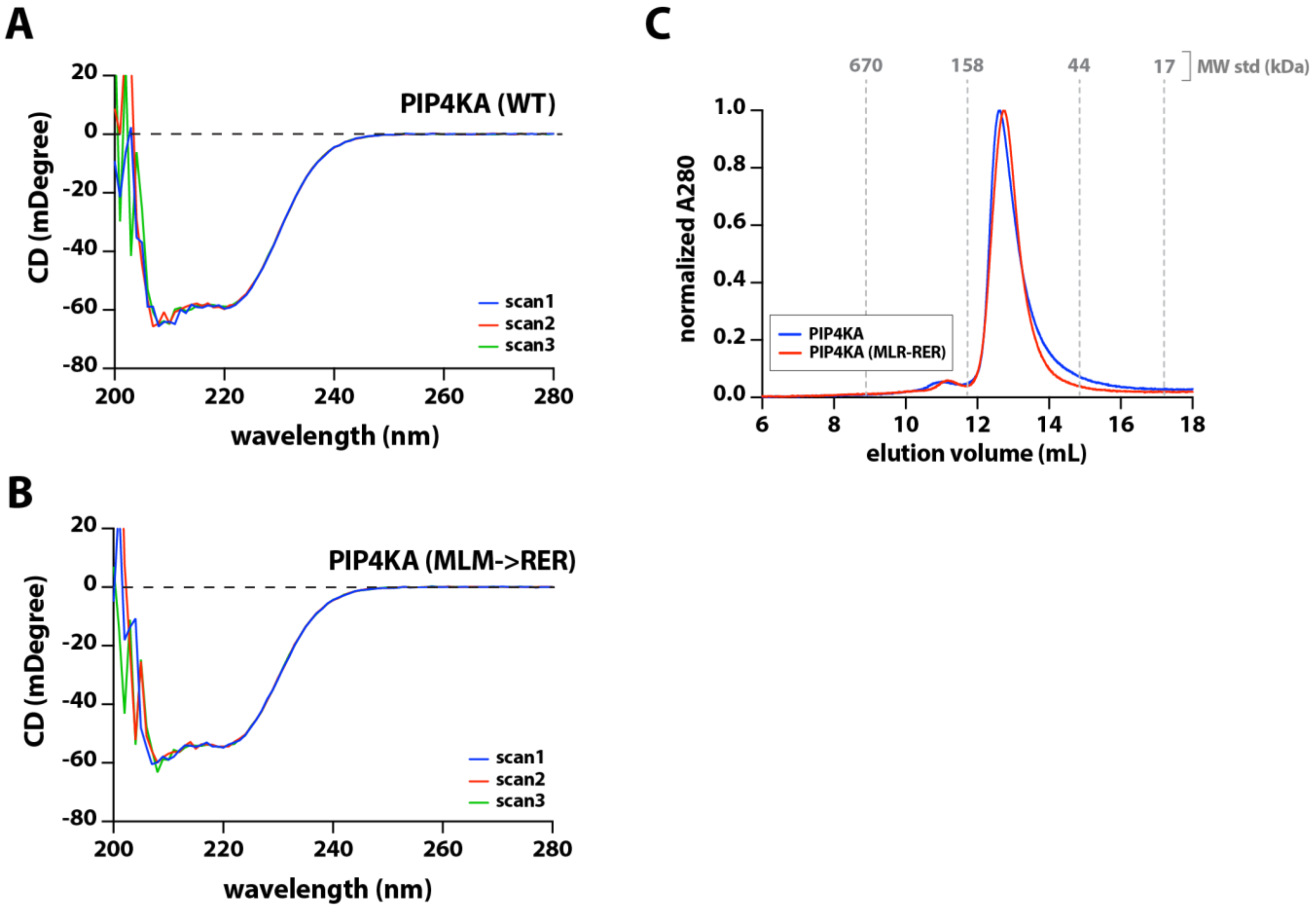


**Supplemental Figure 7**

**PIP4KA, wild type and mutant, are structural indistinguishable based on circular dichroism (CD) and size exclusion chromatography (SEC)**

**(A-B)** Circular dichroism indicates that PIP4KA, wild type and M60R, L61E, M62R mutant, have indistinguishable secondary structure in solution. Measurements were made in the presence of 15 µM PIP4KA. Plots show overlay of 3 scans. **(C)** PIP4KA, wild type and M60R, L61E, M62R mutant, have similar size exclusion chromatography elution profiles measured on a Superdex200 column. A volume of 0.5 mL of 7.5 µM PIP4K was injected for each SEC run. Elution profile of BioRad molecular weight standards is shown in grey with sizes in kDa.

### Supplemental Figure 8

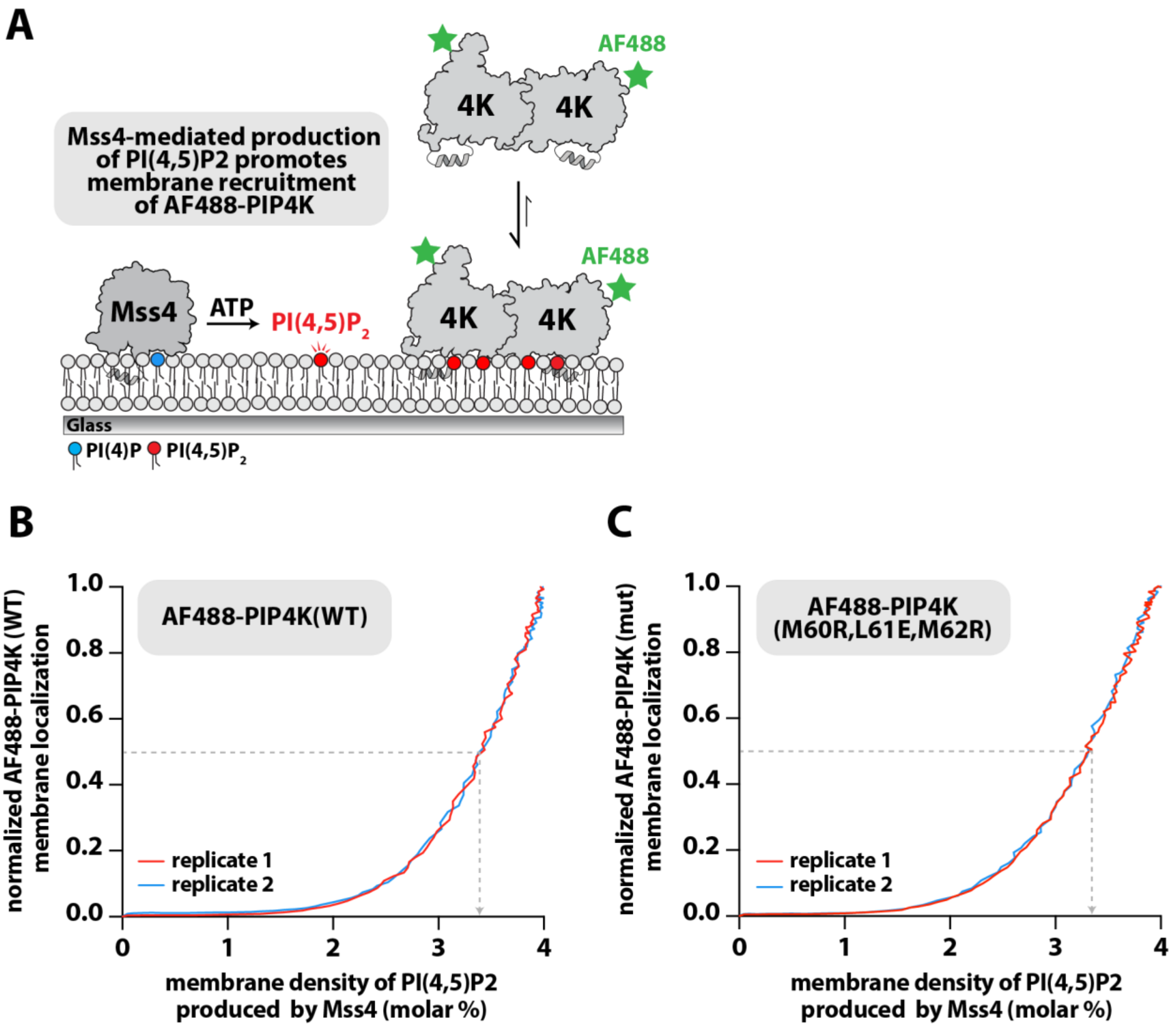


**Supplemental Figure 8**

**PIP4KA, wild type and mutant, bind cooperatively to PI(4,5)$P_2$ membranes**

**(A)** Cartoon schematic showing yeast Mss4-mediated phosphorylation of PI(4)P to generate PI(4,5)$P_2$ on supported lipid bilayers. In the absence of PI(4,5)$P_2$, PIP4K is predominantly in solution. Production of PI(4,5)$P_2$ drives membrane localization of PIP4K. We previously showed that Mss4 does not interact with PIP4K (Wills et al., 2023). **(B-C)** Mss4-mediated production of PI(4,5)$P_2$ promotes cooperative membrane recruitment of AF488-PIP4K. Reactions contain 10 nM Mss4, 50 nM AF488-PIP4KA (WT or M60R, L61E, M62R mutant), 20 nM AF647-PLCδ PH domain. The intensity of membrane localized AF488-PIP4K was normalized and plotted against the density of PI(4,5)$P_2$ generated by Mss4. The dashed grey line represents the molar density of PI(4,5)$P_2$ that resulted in 50% of maximum **(B)** AF488-PIP4K (3.4% $PIP_2$) or **(C)** AF488-PIP4K mutant (3.3% $PIP_2$) membrane density. Initial membrane composition: 4% PI(4)P, 96% DOPC.

### Supplemental Figure 9

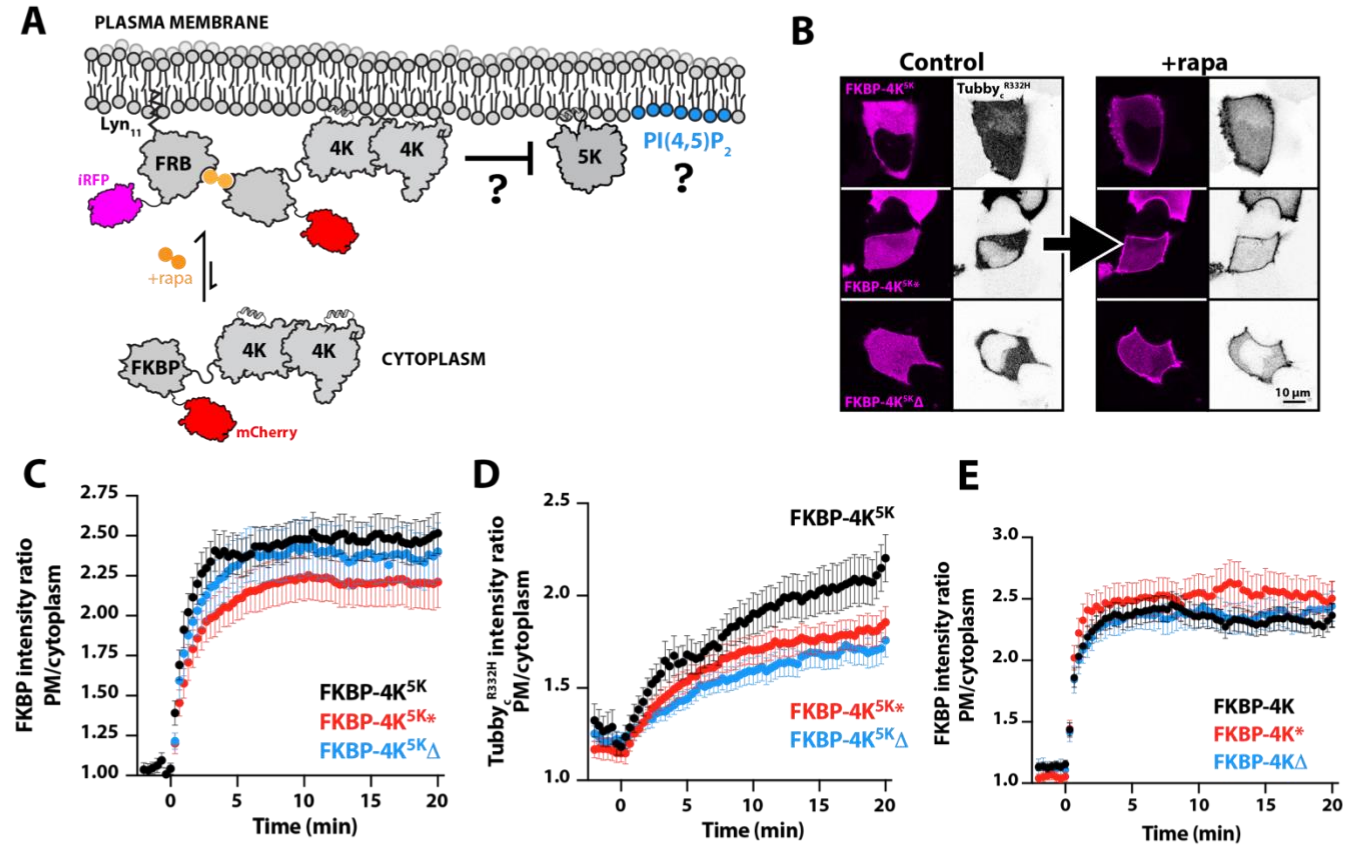


**Supplemental Figure 9**

**Rapamycin induced heterodimerization drives plasma membrane localization FKBP-PIP4KA**

**(A)** Cartoon illustrating the mechanism of recruiting cytosolic mCherry-FKBP-PIP4KA to the plasma membrane in the presence of rapamycin and membrane anchored $Lyn_{11}$-FRB-iRFP. **(B)** Representative confocal sections of HEK293A cells expressing low affinity $PI(4,5)P_2$ biosensor $Tubby_c^{R332H}$-mCherry, $Lyn_{11}$-FRB-iRFP (not shown) and either mCherry-tagged FKBP-$4K^{5K}$ (where 5K indicates the A371E substrate swap mutation conferring PI4P 5-K activity), -$4K^{5K*}$ or $4K^{5K}\Delta$. Images are shown before or 20 min after the addition of 1 μM rapamycin to induce FRB-FKBP dimerization and PM recruitment of the 4K constructs. **(C)** FKBP-$4K^{5K}$ constructs containing the A371E substrate swap mutation display similar rapamycin induced plasma membrane localization. Quantification of time lapse experiments in (B), showing the fluorescence ratio at the plasma membrane relative to cytoplasm for FKBP-$4K^{5K}$ constructs. Data are means ± s.e.m. of 34-49 cells from 3-5 experiments. **(D)** Quantification of time lapse experiments in (B), showing the fluorescence ratio at the plasma membrane relative to cytoplasm $Tubby_c^{R332H}$. Data are means ± s.e.m. of 34-49 cells from 3-5 experiments. **(E)** Quantification of time lapse experiments for the experiment from Figure 6E, showing the fluorescence ratio at the plasma membrane relative to cytoplasm for FKBP-4K constructs. Data are means ± s.e.m. of 36-78 cells from 3-7 experiments.

#### SUPPLEMENTAL MOVIE LEGEND

**Movie S1**
**Visualization of PIP5KB membrane-mediated dimerization by smFRET**
AF647-PIP5KB acceptor emission visualized by smFRET in the presence of AF555-PIP5KB (donor, not shown) on a supported lipid bilayer. Movie contains two example tracks from separate membrane locations merged into a single movie. Data collected in the presence of 5 nM AF555-PIP5KB (donor) and 1 nM AF647-PIP5KB (acceptor). Membrane composition: 4% PI(4,5)$P_2$, 96% DOPC. Movie plays at 10 fps. Data was collected with 52 ms time intervals (19 fps). Spot detection (yellow) and trajectories (purple) overlaid. Scale bar equals 2 μm.

### PLASMID INVENTORY

| Plasmid (main text name) | Vector | Gene sequence, insert, open reading frame, etc. | Reference |
|---|---|---|---|
| PLCδ | pETM | his6-TEV-SUMO3-GGGGG-PLCδ PH domain (11-140aa) | 1 |
| LactC2 | pETM | his6-TEV-SUMO3-GGGGG-LactC2 (271-427aa) | 2 |
| PIP5KB | FastBac | his6-MBP-TEV-GGGGG-PIP5KB (1-421aa) | 3 |
| PIP5KB (D51R) | FastBac | his6-MBP-TEV-GGGGG- PIP5KB (1-421aa) D51R | 3 |
| PIP5KA | FastBac | his6-MBP-N10-TEV-GGGGG-mPIP5K1A (1-546aa) | 3 |
| PIP5KA (D92R) | FastBac | his6-MBP-N10-TEV-GGGGG-mPIP5K1A (1-546aa) D92R | 3 |
| PIP5KB FRET | FastBac | his6-MBP-N10-TEV-GGGG-PIP5KB (1-421aa; C110S, P163C, C411S) | This study |
| PIP5KB FRET (D51R) | FastBac | his6-MBP-N10-TEV-GGGG-PIP5KB (1-421aa; D51R, C110S, P163C, C411S) | This study |
| PIP5KB FRET (D266K) | FastBac | his6-MBP-N10-TEV-GGGG-PIP5KB (1-421aa; D266K, C110S, P163C, C411S) | This study |
| PIP4KA | pETM | his6-TEV-SUMO3-GGGGG-PIP4K2A | 4 |
| PIP4KA (R104D) | pETM | his6-TEV-SUMO3-GGGGG-PIP4K2A (R104D) | This study |
| PIP4KA (M60R,L61E,M62R) | pETM | his6-TEV-SUMO3-GGGGG-PIP4K2A (M60R,L61E,M62R) | This study |
| Mss4 | pETM | his6-MBP-N10-TEV-GGGGG-Mss4 (376-779aa) | 4 |
| EGFP-PIP5K1A | pEGFP-C1 | EGFP:PIP5K1A | 4 |
| TagBFP2 | pTagBFP2-C1 | mTagBFP2 | 4 |
| TagBFP2-4K | pTagBFP2-C1 | mTagBFP2:PIP4K2A | 5 |
| TagBFP2-4K* | pTagBFP2-C1 | mTagBFP2:PIP4K2A$^{M60R,L61E,M62R}$ | This study |
| TagBFP2-4K$^{\Delta}$ | pTagBFP2-C1 | mTagBFP2:PIP4K2A$^{\Delta 54\text{-}62 \rightarrow GGSGG}$ | This study |
| FKBP-4KA | pmCherry-C1 | mCherry:FKBP1A(3-108):[GGSA]$_4$GG:PIP4K2A | 5 |
| FKBP-4K* | pmCherry-C1 | mCherry:FKBP1A(3-108):[GGSA]$_4$GG:PIP4K2A$^{M60R,L61E,M62R}$ | This study |
| FKBP-4K$^{\Delta}$ | pmCherry-C1 | mCherry:FKBP1A(3-108):[GGSA]$_4$GG:PIP4K2A$^{\Delta 54\text{-}62 \rightarrow GGSGG}$ | This study |
| FKBP-4KA$^{5K}$ | pmCherry-C1 | mCherry:FKBP1A(3-108):[GGSA]$_4$GG:PIP4K2A$^{A371E}$ | This study |
| FKBP-4K$^{5K*}$ | pmCherry-C1 | mCherry:FKBP1A(3-108):[GGSA]$_4$GG:$^{PIP4K2AM60R,L61E,M62R, A371E}$ | This study |
| FKBP-4K$^{5K\Delta}$ | pmCherry-C1 | mCherry:FKBP1A(3-108):[GGSA]$_4$GG:PIP4K2A$^{\Delta 54\text{-}62 \rightarrow GGSGG, A371E}$ | This study |
| Lyn$_{11}$-FRB-iRFP | piRFP-N1 | LYN(1-11):MTOR(2021-2113):iRFP | 6 |
| Tubby$_C$-EGFP | pEGFP-N1 | *Mus musculus* Tub(243-505):EGFP | 7 |
| Tubby$_C^{R332H}$-EGFP | pEGFP-N1 | *Mus musculus* Tub(243-505)(R332H):mCherry | 7 |
| Tubby$_C$-mCherry | pmCherry-N1 | *Mus musculus* Tub(243-505):mCherry | 7 |
| Tubby$_C^{R332H}$-mCherry | pmCherry-N1 | *Mus musculus* Tub(243-505)(R332H):EGFP | 7 |
| Lyn$_{11}^{C3S}$-EGFP | pEGFP-N1 | LYN(1-11)(C3S):EGFP | 4 |
| Myr-mCh-4K | pEGFP-C1 | EGFP:LYN(1-11)(C3S):PIP4K2A | 4 |
| Myr-mCh-4K* | pEGFP-C1 | EGFP:LYN(1-11)(C3S): PIP4K2A$^{M60R,L61E,M62R}$ | This study |
| Myr-mCh-4K$^{\Delta}$ | pEGFP-C1 | EGFP:LYN(1-11)(C3S): PIP4K2A$^{\Delta 54\text{-}62 \rightarrow GGSGG}$ | This study |

.
